# Neuromodulation of the endbulb of Held synapse in the cochlear nucleus

**DOI:** 10.1101/2025.02.12.637882

**Authors:** Maria Groshkova, Theocharis Alvanos, Yumeng Qi, Fangfang Wang, Carolin Wichmann, Yunfeng Hua, Tobias Moser

## Abstract

Synapses vary greatly in synaptic strength and plasticity, even within the same circuitry or set of pre- and postsynaptic neurons. Neuromodulation is a candidate mechanism to explain some of this variability. Neuromodulators such as monoamines can differentially regulate presynaptic function as well as neuronal excitability. Variability is found also for the large calyceal synapses of the auditory pathway that are endowed with high synaptic vesicle (SV) release probability (P_vr_) and large postsynaptic currents enabling reliable and temporally precise transmission of auditory information. Here we investigated whether the calyceal endbulb of Held synapse formed by auditory nerve fibers onto bushy cells (BCs) in the anteroventral cochlear nucleus (AVCN) is modulated by norepinephrine (NE) and serotonin (5-HT).

Using electron microscopy (EM) of the cochlear nucleus we found evidence for putative monoaminergic varicosities in both ventral and dorsal divisions. Immunostaining for vesicular 5-HT and NE transporters revealed NE-containing and 5-HT-containing varicosities in the AVCN, juxtaposed to both endbulbs and BCs. Furthermore, we detected immunofluorescence for 5-HT_1B_, 5-HT_4_, 5-H_7_ receptors (R) and α_2C_-adrenergic receptors (AR) in BCs. We used voltage-clamp recordings from mouse BCs in order to uncover potential presynaptic effects of neuromodulation, which revealed an increase in frequency of miniature excitatory postsynaptic currents (mEPSCs) upon application of NE but not 5-HT. Evoked synaptic transmission was unaffected by the application of either NE or 5-HT. Likewise, while studying the biophysical properties of the BCs, we did not observe effects of NE or 5-HT on low-voltage-activated K^+^ (K^+^_LVA_) and hyperpolarization-activated mixed cation (HCN) channels during application. In summary, we report evidence for the presence of monoaminergic innervation in the cochlear nucleus and for subtle functional NE-neuromodulation at the endbulb of Held synapse.

## 1 Introduction

The endbulb synapse is characterized by synchronous release from dozens of AZs for producing strong EPSCs for reliable and temporally precise transmission (Wang and Manis, 2005; Wang et al., 2010). In this way this synapse enables a faithful representation of sound’s temporal properties into the central auditory system and allow the performance of tasks such as encoding the onset of sound (Rhode, 2008) and sound localization (Kuenzel, 2019). A recent serial block face scanning EM (SBEM) study in the mouse AVCN described endbulbs, variable in size converging onto the same globular BC (Spirou et al., 2023). Based on compartmental modelling the study suggested that variability in synaptic weights of different inputs contributed to improving the temporal precision of the neural code, leaving the AVCN, that had been reported (Joris et al., 1994).

Aside from strong glutamatergic axosomatic input from the endbulbs formed by the spiral ganglion neurons (SGNs), temporally precise firing of BCs capitalizes on I) fast rapidly desensitizing AMPA receptors (Gardner et al., 1999) and II) short membrane time-constant (τ_m_), which limits the time window of integration (Kuenzel, 2019). The fast τ_m_ corresponds to low input resistance, reflecting the presence of a low-voltage-activated potassium conductance (g_KL_) in BCs. The low-voltage activated potassium channels (K^+^_LVA_) mediating g_KL_ activate near the resting potential, producing large hyperpolarizing currents that prevent BCs from repetitive firing (Cao and Oertel, 2010). Furthermore, g_KL_ makes BCs sensitive to the rate of depolarization, by suppressing slow membrane depolarizations. An opposing hyperpolarization-activated cation conductance (g_h_), mediated by hyperpolarization and cyclic nucleotide activated (HCN) channels, is activated when BCs are exposed to strong hyperpolarizing currents. In BC there is evidence for fast gating HCN1 channels (Oertel et al., 2008). This conductance shortens the refractory period of the BCs, enabling them to receive and respond to high frequency repetitive stimulation from SGNs (Cao et al., 2007). Hence, both g_KL_ and g_h_ define the electrical properties of BC and consequently the excitability of the cells.

While being tuned to reliable and precise transmission of auditory information, evidence for neuromodulation in the cochlear nucleus (CN) has been presented. Neuromodulators are molecules that can modify synaptic transmission and the excitability of the postsynaptic cell by affecting the properties of synaptic vesicle (SV) release, neurotransmitter receptors, or voltage-gated ion channels (Brzosko et al., 2019; Özçete et al., 2024). This effect is achieved through the activity of G protein-coupled receptors (GPCRs) resulting in second messenger cascades of signaling. In fact, a number of G-protein-modulated signal pathways have been shown to influence neurotransmitter release (de Jong and Verhage, 2009). Evidence for modulation was shown for the auditory pathway where calyceal synapses undergo GABA_B_ modulation (Takahashi et al., 1998; Brenowitz and Trussell, 2001). Modulation of presynaptic Ca^2+^ currents (Takahashi et al., 1998; Kimura et al., 2003) is probably the most effective way to influence neurotransmitter release due to the supralinear relationship between release and intracellular [Ca^2+^]. However, several modulators act via G_q_-signaling coupled to the PLC-diacylglycerol (DAG) pathway for which the priming protein Munc13 is a prominent target. Although it has been known for decades that phorbol ester, a mimic of DAG, increases synaptic strength by factors of 2 to 6 (e.g. Hori et al., 1999; Lou et al., 2005; Lee et al., 2013), little attention has been paid so far to the possibility of neuromodulation via this route. Furthermore, G_s_-signaling has been described for the calyx of Held that increased both readily releasable pool and the release probability at the calyx of Held (Sakaba and Neher, 2001; Kaneko and Takahashi, 2004). Candidate neuromodulators of the CN include the monoamines dopamine, 5-HT and NE. Monoamine modulators are mainly secreted in targeted areas in a process called volume transmission (VT) (Agnati et al., 1986; Fuxe and Borroto-Escuela, 2016). In this process the neurotransmitter molecules diffuse through the extracellular fluid and interact with their respective neuromodulator receptors. The projections of neuromodulators-releasing neurons are known to form swellings along their length known as varicosities where the secretion is known to occur from (Séguéla et al., 1990; Descarries et al., 1996; Gianni and Pasqualetti, 2023). These typically unmyelinated swellings contain clear and dense core vesicles and most often lack classical synaptic specializations (Séguéla et al., 1990; Descarries and Mechawar, 2000; Liu et al., 2018). The concentration gradient that results from volume transmission seems to form domains of affinity, corresponding to a non-random compartmental placement of respective modulator receptors in the postsynaptic cells (Özçete et al., 2024).

Noradrenergic synaptic transmission takes part in processes such as arousal, attention, cognition, fear conditioning and memory formation (Groch et al., 2011). NE acts through two broad families of receptors, namely α-and β-adrenergic receptors (ARs). The α-AR family is subdivided to α_1_-ARs, coupled with G_q_ proteins, activating PLC (Wu et al., 1992), leading to an increase of IP_3_ and intracellular Ca^2+^; and α_2_-ARs, acting through G_i/o_ proteins, thus lowering cAMP levels by inhibiting AC. Particularly, α_2_-AR signaling inhibited HCN currents in neurons in the prefrontal cortex (Carr et al., 2007; Wang et al., 2007; Zhang et al., 2013). Furthermore, activation of α_2_-ARs at climbing fibers in the cerebellum decreased P_vr_ and subsequently modulates short-term associative plasticity of the parallel fiber to Purkinje cell synapse (Carey and Regehr, 2009). NE differentially modulated short-term depression of evoked inhibitory transmission at layers I versus layers II/III in the auditory cortex at 20 Hz stimulation frequency (Salgado et al., 2011). While its effect on layer I was to alter the depressive pattern to a facilitating one, in layers II/III the depression was even more prominent after NE application. Additionally, one of the early findings on presynaptic neuromodulation was the inhibitory effect of NE on the release from cholinergic projections in the ileum longitudinal muscle strip of guinea pigs, mediated by α_2_-ARs (Paton and Vizi, 1969). In stem-cell-derived human neurons, the effect of α_2_-AR activation resulted in cAMP reduction and synapsin-1 de-phosphorylation. This augmented the reserve pool of SVs, complementing the 5HT_7_R effect and allowing for bi-directional control of SV replenishment (Patzke et al., 2019). The β-AR family consists of 3 subgroups – β_1_-ARs, β_2_-ARs, β_3_-ARs all acting through G_s_ proteins and elevating cAMP levels (Sibley and Lefkowitz, 1987). β-adrenergic signaling has been linked to inactivation of dendritic Kv1.1 resulting in increased dendritic excitability (Liu et al., 2017). Moreover, noradrenergic modulation was implicated in tuning spike timing dependent plasticity in the visual cortex through activation of AC and PLC cascades (Seol et al., 2007; Huang et al., 2012).

5-HT is synthesized by neurons in the Raphe nuclei enabling serotonergic signaling involved in various physiological processes, including sleep, social behavior, sexual activity, learning and memory, pain, feeding (Bockaert et al., 2006). Serotonergic inputs into neural networks operate via at least 15 structurally and pharmacologically different receptors (Hoyer et al., 1994) and trigger various responses. Among them are hyperpolarization-induced decrease in neural firing rate, stimulation of phospholipase C (PLC) via G_q_ proteins, initiating intracellular Ca^2+^ and diacylglycerol signaling, and activation or inactivation (G_s_ or G_i/o_) of adenylate cyclase (AC) to regulate cyclic AMP (cAMP) levels (Filip and Bader, 2009). Within the dorsal cochlear nucleus (DCN), serotonin enhanced the excitability of fusiform principal cells in the through the activation of 5-HT receptors 5-HT_2A_/_2C_R and 5-HT_7_R via HCN current augmentation (Tang and Trussell, 2015). Furthermore, 5-HT_7_R activation upregulated the phosphorylation of synapsin-1 and the consequent decrease SV replenishment in stem-cell-derived human neurons (Patzke et al., 2019). Additionally, the 5-HT_1B_R, expressed presynaptically, reduced GABA_A_ signaling in the inferior colliculus in the auditory midbrain and thus facilitated higher spiking rates of inferior colliculus neurons (Hurley et al., 2008). Serotonin facilitated long-term depression of glutamatergic transmission at the medium spiny neurons in the Nucleus accumbens by activating 5-HT_1B_Rs (Huang et al., 2013). In the hippocampal dentate gyrus of rats the 5-HT_4_R had an inhibitory effect on long-term potentiation (Kulla and Manahan-Vaughan, 2002).

The presence of noradrenergic and serotonergic innervation of the CN of rat and cat has been indicated through biochemical and fluorescence based essays (Kromer and Moore, 1976; Klepper and Herbert, 1991; Cransac et al., 1995; Thompson and Thompson, 2001). However, a detailed analysis of neuromodulation in the CN remained to be performed. Here, we used serial block-face electron microscopy (SBEM) of the CN and found varicose neurites indicative of monoaminergic innervation. We then focused on studying neuromodulation of the mouse endbulb synapse of SGNs and bushy cells in the AVCN by NE and 5-HT. Immunohistochemistry revealed the presence the norepinephrine and serotonin transporters NET and SERT near the endbulb synapse as well as the expression of α_2C_-AR, 5-HT_1B_R, 5-HT_4_R and 5-HT_7_R receptors in/near BCs supporting the hypothesis of serotonergic and adrenergic modulation in the AVCN. Finally, we performed whole-cell patch-clamp recordings from BC to evaluate the effects of administered NE and 5-HT on the spontaneous and evoked synaptic transmission as well as on the electrical properties of BCs.

## 2 Materials and methods

### 2.1 Animals

Mice from the wild-type substrain C57BL/6N were used for the electrophysiology and immunostaining experiments. They were obtained from the colony maintained at Max Planck Institute for Multidisciplinary Sciences, Faßberg Campus, Göttingen. Male and female mice, of ages ranging between P14 and P20, were sacrificed by decapitation for then dissecting the brain and preparing acute brainstem slices. Mice aged p15 to p22, as well as a p42 mouse, were used in our immunohistochemical assays. The use of animals complied with national animal care guidelines in the registered facility 33.23-42508-066-§11. For the electron microscopy experiments an 8-week-old C57BL/6J mouse was used to make VCN sample and two 7-week-old CBA/Ca mice were used to make DCN samples. The 8-week-old C57BL/6J mouse was purchased from Shanghai Jihui Laboratory Animal Care Co., Ltd., the two 7-week-old CBA/Ca mice were purchased from Sino-British: SIPPR/BK, Lab. Animal Ltd: (Shanghai, China). And the experiments complied with national animal care guidelines and were approved by the Institutional Authority for Laboratory Animal Care of Shanghai Ninth People’s Hospital (SfH9H-2020-A65-1).

### 2.2 SBEM sample preparation

The mice were anesthetized with 2% isoflurane inhalation before successive transcardial perfusions of 15 mL sodium cacodylate buffer (0.15 M) and 30 mL mixed fixative solution containing 2% paraformaldehyde and 2.5% glutaraldehyde (buffered by 0.08 M sodium cacodylate, pH = 7.4). After decapitation, the brains were exposed by removing the skull and post-fixed by immersion in the same fixative at 4°C for 24 hours. Then, the specimen was transferred to sodium cacodylate buffer (0.15 M) in a Petri dish (on ice), and the brainstem was exposed by carefully removing the cerebellum under a dissecting microscope. The CN samples were harvested from the brainstem by cutting with a scalpel blade.

The en bloc staining for SBEM was performed following the previously published protocol (Hua et al., 2015, 2022) with minor modifications. In brief, the CN samples were washed twice in 0.15 M cacodylate (pH 7.4) for 30 min each and sequentially immersed in 2% OsO4 (Ted Pella), 2.5% ferrocyanide (Sigma), and again 2% OsO4 at room temperature (RT) for 2, 1.5, and 1 hours, respectively, without intermediate washing step. All staining solutions were buffered with 0.15 M cacodylate (pH 7.4). After being washed sequentially in 0.15 M cacodylate and nanopore-filtered water for 30 min each, the samples were incubated at RT in 1% thiocarbonhydrazide (aqueous solution) for 1 hour and further stained with 2% OsO4 aqueous solution for 2 hours, 1% uranium acetate at 4°C for 8 hours and at 50°C for 2 hours, as well as 0.03 M lead aspartate solution (pH 5.0 adjusted by KOH, Electron Microscopy Sciences) at 50°C for 2 hours. Between steps, double rinses in nanopore-filtered water for 30 min each were performed.

For resin embedding, the samples were dehydrated through a graded acetone series (50%, 75%, 90%, for 30 min each at 4°C) into pure acetone (3 × 100%, 45 min at RT), followed by infiltration with 1:1 mixtures of acetone and Spurr’s resin monomer (4.1 g ERL 4221, 0.95 g D.E.R^TM^ 736, 5.9 g NSA and 1% 2-Dimethylaminoethanol, DMAE; Sigma-Aldrich) at RT for 8 hours on a rotator. Infiltrated samples were then incubated in pure resin for 8∼12 hours before being placed in embedding molds (Polyscience, Germany) and incubated in a pre-warmed oven (70°C) for 72 hours.

### 2.3 SBEM imaging

The embedded samples of AVCN and DCN were trimmed to a block face of 600 μm × 600 μm and coated with thin-layer gold (thickness of 30 nm) by a high-vacuum coating device (ACE600, Leica, Germany). It was imaged using a field-emission SEM (Gemini300, Zeiss) equipped with an in-chamber ultramicrotome (3ViewXP, Gatan). Focal charge compensation was set to 100% with a high vacuum chamber pressure of 2.8 × 103 mbar. Serial images were acquired in a stitching mode at 12 nm or 15 nm pixel size and nominal cutting thickness of 35 or 50 nm; incident beam energy 2 keV; dwell time 1.5 μs.

### 2.4 Electrophysiology solutions

Dissections were performed in an ice-cold cutting solution containing (in mM): 50 NaCl, 26 NaHCO_3_, 1.25 NaH_2_PO_4_H_2_O, 2.5 KCl, 20 glucose, 0.2 CaCl_2_, 6mM MgCl_2_, 0.7 Na L-ascorbate, 2 Na pyruvate, 3 myo-inositol, 3 Na L-lactate, 120 sucrose adjusted to pH 7.3–7.4. Recordings were performed in solutions based on artificial cerebrospinal fluid (aCSF), containing (in mM): 125 NaCl, 26 NaHCO_3_, 1.25 NaH_2_PO_4_H_2_O, 2.5 KCl, 13 glucose, 2 CaCl_2_, 1 MgCl_2_, 0.7 Na L-ascorbate, 2 Na pyruvate, 3 myo-inositol, 3 Na L-lactate adjusted to pH 7.3–7.4. aCSF was supplemented with 10 μM bicuculline, a competitive antagonist of GABA_A_ receptors and 2µM strychnine to inhibit postsynaptic glycine receptors (control solution). The test solution additionally contained either 100 μM NE or 10 μM 5-HT. All solutions were continuously aerated with carbogen (95% O_2_,5%CO_2_).

For the HCN and K^+^_LVA_ experiments the control and test solutions additionally contained 1 µM tetrodotoxin (TTX, Alomone labs) blocking voltage-sensitive sodium currents, 0.25 mM CdCl_2_ blocking the voltage-sensitive Ca^2+^ currents; 10 µM 6,7-dinitroquinoxaline-2,3-dione (DNQX) to block excitatory postsynaptic currents (EPSCs). To measure HCN currents additional 25 nM α-dendrotoxin (α -DTX, Alomone labs) was added to block K^+^_LVA_. For the measurement of K^+^_LVA_ currents we included 10 µM ZD7288 to block HCN channels.

During the fiber stimulation experiments the control solution was supplemented with 1 mM kynurenic acid sodium salt (abcam Biochemicals, Cambridge, UK), a low-affinity AMPAR antagonist, to prevent receptor saturation/desensitization.

The pipette solution contained (in mM): 115 K-gluconate, 8 EGTA, 10 HEPES, 4 Mg-ATP, 0.3 Na-GTP, 10 Na_2_Phosphocreatine, 4,5 MgCl_2_, 10 NaCl, pH 7.3, 317 mOsm. Additionally, the fluorescent dye Alexa 568 (34 µM, Invitrogen) was added to the intracellular solution to assist identification of the cell type which we observed using a HXP 120184 mercury lamp, with a FITC filter set (Semrock). For the fiber stimulation experiments we added 1 N-(2,6-dimethylphenyl carbamoylmethyl) triethylammonium chloride (QX-314; Alomone Labs, Jerusalem, Israel) to the intracellular solution to block sodium channels. The source of the above listed chemicals was Merck, Germany unless stated differently.

### 2.5 Slice preparation for electrophysiology

Following sacrifice of mice, their brains were immediately immersed in ice-cold cutting solution. A midsagittal cut was performed on the brain and the hindbrain was severed from the forebrain. This was followed by gluing the hindbrain to the stage of the vibratome (VT 1200S, Leica, Wetzlar, Germany). To obtain parasagittal brainstem slices a 0.02 mm/s advancing rate of the cutting blade and a vibration amplitude of 1.5 mm were used. 150μm thick parasagittal slices of the CN were cut and incubated for 30 minutes in aCSF at 35°C. Dissection materials: forceps, small surgical scissors, surgical scissors, wax stage, dissection blade, glass dropper (all Fine Science Tools, Heidelberg, Germany), cyanoacrylate glue Loctite 401 (Henkel, Germany).

### 2.6 Electrophysiological recordings

Patch pipettes were made from borosilicate glass (Science products, GB150F-8P) and pulled using a P-97 Flaming/Brown micropipette puller (Sutter Instruments). The pipettes had resistances between 2.5 and 4.5 MΩ. The slices were placed in a custom-made stage on an Axioscope 2 FS plus microscope (Zeiss, Jena, Germany) and held in place with a custom-made stainless-steel harp with glued nylon strings. The slices were visualized with a 40x water-immersion objective, using differential interference contrast illumination. Patch clamp recordings were made from BC of the AVCN using an EPC10 USB double patch-clamp amplifier, which was controlled by the PatchMaster software (HEKA Elektronik). The sampling interval was 25μs and data were filtered at 7.3kHz. A liquid junction potential correction of 12 mV was performed through the PatchMaster software and cells were voltage clamped at a holding potential of −70mV. Mean series resistance was around 5 MΩ and was compensated up to 70% with a 10µs lag. Recordings, which displayed leak currents beyond −150 pA at −70 mV or a series resistance (R_s_) above 10 MOhm were discarded. All experiments were performed at near physiological temperature (32-35 °C) maintained by constant superfusion (flow rate 3-4ml/min) of aCSF, heated by an inline solution heater (SH-27B174 with TC-324B controller; Warner Instruments, Hamden, CT, USA) and monitored by a thermistor placed between the inflow site and the slice, in the recording chamber.

The recordings of mEPSCs, rheobase and rate of depolarization from cells that compose our control dataset speed were made in the following fashion: after achieving whole-cell configuration while perfusing aCSF supplemented with 10 µM strychnine, we tested for the cell identity. This was done by briefly recording mEPSCs (BC mEPSCs show shorter decay times then stellate cells) as well as current clamp protocols testing the sensitivity to speed of depolarization, the firing patterns of the cells evoked by stepwise current injection (BCs show phasic, stellate cells tonic firing, Fig. 6A_I_). For estimating the rate of depolarization, we documented the slowest rate at which an AP is generated and analyzed the first derivative of the voltage trace before the generation of the AP (Fig. 6A_I_). To estimate the rheobase, we injected currents with different amplitudes and duration and documented the first current value at which an action potential (AP) was generated (Fig. 6B_I_). Once we had determined the cell to be a BC, we perfused the control solution for 1 minute. At 1 min of control solution perfusion, we recorded mEPSC for 1 minute, right after that we performed the above-mentioned current clamp protocols. The recordings from cells exposed to 5-HT or NE were performed in a similar fashion, where, after we determined the cell type to be BC, we perfused the test solution containing 5-HT or NE. At 5 min of perfusion we recorded mEPSCs, followed by the aforementioned current clamp protocols.

The K^+^_LVA_ and HCN channel experiments were performed separately for control, 5-HT and NE conditions after incubation with the respective solution for 5 minutes. In the HCN channel experiments we voltage clamped the cell and changed the voltage in steps of 5 mV from −57 to −112 mV. For the K^+^_LVA_ experiments we used voltage steps of 5 mV from −90 mV to −40 mV. We derived the HCN currents at the plateau at −112 mV and the K^+^_LVA_ currents at −40 mV, then normalized them by the capacitance of the cell in order to derive the current density. As a negative control for the HCN experiments 10 µM of ZD7288 was added to this solution to block HCN currents (Fig. 8A). As a negative control for the K^+^_LVA_ 50 nM α-DTX was added to block K^+^_LVA_ (Fig. 7A).

For studying effects on evoked endbulb transmission, we minimally stimulated presynaptic auditory nerve fibers forming the endbulbs of Held with a monopolar electrode in a patch pipette filled with aCSF, placed at a distance of at least three cell diameters from the cell being recorded. Stimulating currents of 7-24µA were delivered through a stimulus isolator (A360 World Precision Instruments, Sarasota, FL, USA). We performed recordings of regular trains of 35 evoked EPSC (eEPSCs) at 100 Hz and 200 Hz. We perfused the control solution for this experiment for 2 minutes before recording. Next, we perfused the NE or 5-HT containing solutions for 5 minutes before starting to record the second set of data.

### 2.7 Sample preparation for immunohistochemistry

The whole brain was submerged in a 4% formaldehyde (FA, Carl Roth, Karlsruhe, Germany) in PBS (Merck, Germany) for fixation, right after detachment from the skull. The brain was fixed for 24h at 4°C and, thereafter, the solution was changed to 30% PBS-based sucrose solution to dehydrate the sample and prepare it for the next step of cryo-sectioning. The brains were incubated in the sucrose solution for approximately 2 days until they sunk to the bottom of the falcon tube. The cryostat was set to −22°C for the chamber and to −23°C for the stage. Before sectioning, the forebrain and hindbrain were separated and the dehydrated hindbrains were attached to the cutting stage by immersion in the Cryomatrix (ThermoFisher Scientific, Waltham, MA, USA). This was followed by coronal cryo-sectioning in a caudal to rostral direction. The anatomical landmark for reaching the level of the AVCN is the 7th cranial nerve which appears as an opaque white line traversing the brainstem in a ventrolateral to dorsomedial orientation. Subsequent 30 μm thick cryosections, containing AVCN were collected on electrostatically positively charged microscope slides (ThermoFisher Scientific, MA, USA) to a total amount of 8 slices of each ACVN. The collected slices were kept at −20°C before fixation and staining. Before staining each slice was incubated with blocking solution (goat serum dilution buffer, GSDB: 16% normal goat serum, 450 mM NaCl, 0.6% Triton X-100, 20 mM phosphate buffer) for 1 h.

### 2.8 Staining and microscopy

All antibodies (Ab) were diluted in GSDB. For staining of monoamine transporters, antibodies against norepinephrine (NET) and serotonin (SERT) transporters were diluted 1:200 and for counterstaining the presynaptic marker vGlut1 guinea-pig-anti-vGlut1 was diluted 1:2000. In the case of the monoamine receptor stainings, primary antibody combinations routinely contained guinea-pig-anti-vGlut1 Ab (1:1000 or 1:2000, Synaptic Systems GmbH, Göttingen, Germany), either chicken anti-Homer1 Ab (1:500, Synaptic Systems GmbH, Göttingen, Germany), labeling the excitatory postsynaptic density inside of the principal cell or mouse Gephyrin (1:500, Synaptic Systems GmbH, Göttingen, Germany), labeling the postsynaptic density of inhibitory synapses. In order to label for the various receptors, a relevant rabbit Ab (1:200 or 1:500) was included used. In parallel, a negative control staining with the same components and with additional blocking peptide for each of the tested receptors was performed. The concentration of the blocking peptide was 500 higher than that of the antibody in the final solution. Before applying to the tissue, both solutions were kept at room temperature and centrifuged at 2000 g for an hour. All the slices were incubated with the primary antibodies overnight at 4°C.

This was followed by washing 3×10 mins first with washing buffer (20 mM Phosphate buffer, 0.3% Triton X-100, 0.45 M NaCl) and then 3×10 mins with PBS. The slides were dried off of PBS and secondary antibodies goat-anti-rabbit Alexa fluor 488, goat-anti-chicken/mouse Alexa fluor 568, guinea-pig-anti-mouse Alexa fluor 633/647 were added at a dilution of 1:200. The incubation with secondary Abs lasted 2 hours and was followed by washing 3×10 minutes with wash buffer and 3×10 minutes with PBS. Next, the slides were washed in 5 mM phosphate buffer for 5 minutes and mounted with 50 µl of fluorescence mounting medium based on Mowiol 4-88 (Carl Roth, Karlsruhe, Germany) and covered with a thin glass coverslip. The slides were kept at 4°C until imaging. The list of all antibodies used can be found on Tables 1 and 2.

**Table 1.** Primary antibodies used in the immunolabelling experiments.

| Primary Antibodies | Source |
| --- | --- |
| **mouse-anti-Norepinephrine transporter** | Synaptic Systems GmbH, Göttingen, Germany |
| **rabbit-anti-Serotonin transporter** | Synaptic Systems GmbH, Göttingen, Germany |
| **guinea-pig-anti-vGlut1** | Synaptic Systems GmbH, Göttingen, Germany |
| **Chicken-anti-Homer1** | Synaptic Systems GmbH, Göttingen, Germany |
| **Mouse-anti-Gephyrin** | Synaptic Systems GmbH, Göttingen, Germany |
| **rabbit-anti-5-HT<sub>1A</sub> receptor +blocking peptide** | Alomone Labs, Israel |
| **rabbit-anti-5-HT<sub>1B</sub> receptor +blocking peptide** | Alomone Labs, Israel |
| **rabbit-anti-5-HT<sub>2A</sub> receptor +blocking peptide** | Alomone Labs, Israel |
| **rabbit-anti-5-HT<sub>2B</sub> receptor +blocking peptide** | Alomone Labs, Israel |
| **rabbit-anti-5-HT<sub>4</sub> receptor +blocking peptide** | Alomone Labs, Israel |
| **rabbit-anti-5-HT<sub>5B</sub> receptor +blocking peptide** | Alomone Labs, Israel |
| **rabbit-anti-5-HT<sub>7</sub> receptor, +blocking peptide** | Alomone Labs, Israel |
| **rabbit-anti-$\alpha_{1B}$-AR, +blocking peptide** | Alomone Labs, Israel |
| **rabbit-anti-$\alpha_{1D}$-AR, +blocking peptide** | Alomone Labs, Israel |
| **rabbit-anti-$\alpha_{2C}$-AR, +blocking peptide** | Alomone Labs, Israel |
| **rabbit-anti-$\beta_1$-AR, +blocking peptide** | Alomone Labs, Israel |
| **rabbit-anti-$\beta_2$-AR, +blocking peptide** | Alomone Labs, Israel |
| **rabbit-anti-$\beta_3$-AR, +blocking peptide** | Alomone Labs, Israel |

**Table 2.** Secondary antibodies used in the immunolabelling experiments.

| Secondary antibodies |  | Source |
| --- | --- | --- |
| Dilution 1:200 |  |  |
| Fluorophore | Coupled to, Ab |  |
| Alexa fluor 488 | Goat-anti-guinea-pig | MoBiTec (Invitrogen), Göttingen, Germany |
| Alexa fluor 488 | Goat-anti-rabbit | MoBiTec (Invitrogen), Göttingen, Germany |
| Alexa fluor 568 | Goat-anti-guinea-pig | MoBiTec (Invitrogen), Göttingen, Germany |
| Alexa fluor 568 | Goat-anti-rabbit | Thermo Fisher, Waltham, Massachusetts, US |
| Alexa fluor 568 | Goat-anti-chicken | Abcam, Cambridge, UK |
| Alexa fluor 633 | Goat-anti-guinea-pig | ThermoFisher, Waltham, Massachusetts, US |
| Alexa fluor 647 | Goat-anti-guinea-pig | ThermoFisher, Waltham, Massachusetts, US |
| Alexa fluor 647 | Goat-anti-rabbit | MoBiTec (Invitrogen), Göttingen, Germany |
| Alexa fluor 633 | Goat-anti-mouse | ThermoFisher, Waltham, Massachusetts, US |
| Abberior Star 635p | goat-anti-mouse 635p | Abberior, Göttingen, Germany |
| Abberior Star 580 | Goat-anti-guinea-pig | Abberior, Göttingen, Germany |

Two-colour STED images of cryosections stained for monoamine transporters were acquired using a STEDYCON (Abberior Instruments GmbH, Göttingen, Germany) with a 100x oil immersion objective. Images of the monoamine receptor preparations were taken with Zeiss confocal laser scanning microscope 780 and 880 with 40x or 100x oil immersion objectives.

### 2.9 Data analysis

#### 2.9.1 SBEM image processing, neurite reconstruction, and quantification

The dataset was aligned using a self-written MATLAB script based on cross-correlation maximum between consecutive slices. Then, the aligned datasets were split into cubes (1024 × 1024 × 1024 voxels) for viewing and tracing in a browser-based annotation tool, webKNOSSOS as previously described (Gour et al., 2021; Hua et al., 2022). Neurites with associated mitochondria, dence core vesicles and synaptic vesicle clouds, were volume traced by human annotators.

#### 2.9.2 Patch-clamp analysis

Data were analyzed with Igor pro 6.3 software (Wavemetrics) using custom written programs.

#### 2.9.3 Analysis of confocal data

The obtained images were processed with ImageJ. The Z stacks obtained from the monoamine receptor stainings were transformed into maximum intensity projections. Following that, the immunolabelling figures were created with Inkscape.

#### 2.9.4 Statistical analysis

For the fiber stimulation experiments we used consecutive recordings from the same cells: first subjecting them to a control solution, followed by the test recording in 5-HT or NE solution. For all other experiments we pulled the data in the different datasets from different cells. The data were analyzed using Excel and Igor Pro software. The data is presented as mean ± standard error of the mean (SEM). The number of the animals is indicated as N. For two sample comparisons, normality of the distributions (Jarque-Bera test) and equality of variances (F-test) were tested. This was followed by Student’s t-test or Mann-Whitney-Wilcoxon test in case the normality of distributions and/or equality of variances were not met. Significant differences are presented as *p<0.05, **p<0.01, ***p<0.001.

## 3 Results

### 3.1 Electron microscopical correlates of monoaminergic innervation of the cochlear nucleus

Noradrenergic varicosity diameters in the mouse olfactory bulb were reported to be around 0.2 µm in transverse diameter (Horie et al., 2021), while in rat neocortex - in the range of 0.4 to 1.2 µm (Séguéla et al., 1990). Dopaminergic varicosities were reported to be of an average transverse diameter of 0.24 µm (Descarries et al., 1996). Dense core vesicles were observed in 5-HT and NE neurons in addition to clear core SVs (Suzuki et al., 2015; Horie et al., 2021). Hence, we performed scanning electron microscopy on large sample block faces of the en bloc-stained CN tissues of mice. This allowed us to search for varicosities containing dense core vesicles within both AVCN (Fig. 1A, B) and DCN (Fig. 1C, D) subdivisions. Next, we acquired two small EM volumes using SBEM to 3D reconstruct the candidate projections with consecutive varicosities, which contained dense core vesicles (DCV). The swellings contained synaptic vesicles, DCV and sometimes mitochondria and were about 1 µM in transverse diameter (Fig. 1E). In one of our samples (Fig. 1E, below) a putative postsynaptic density was found. Hence, our SBEM data points out the presence of varicosity-like neurites in the CN and future investigations on larger EM volumes of AVCN will be needed to comprehensively characterize this type of neurite.

**Figure 1.**
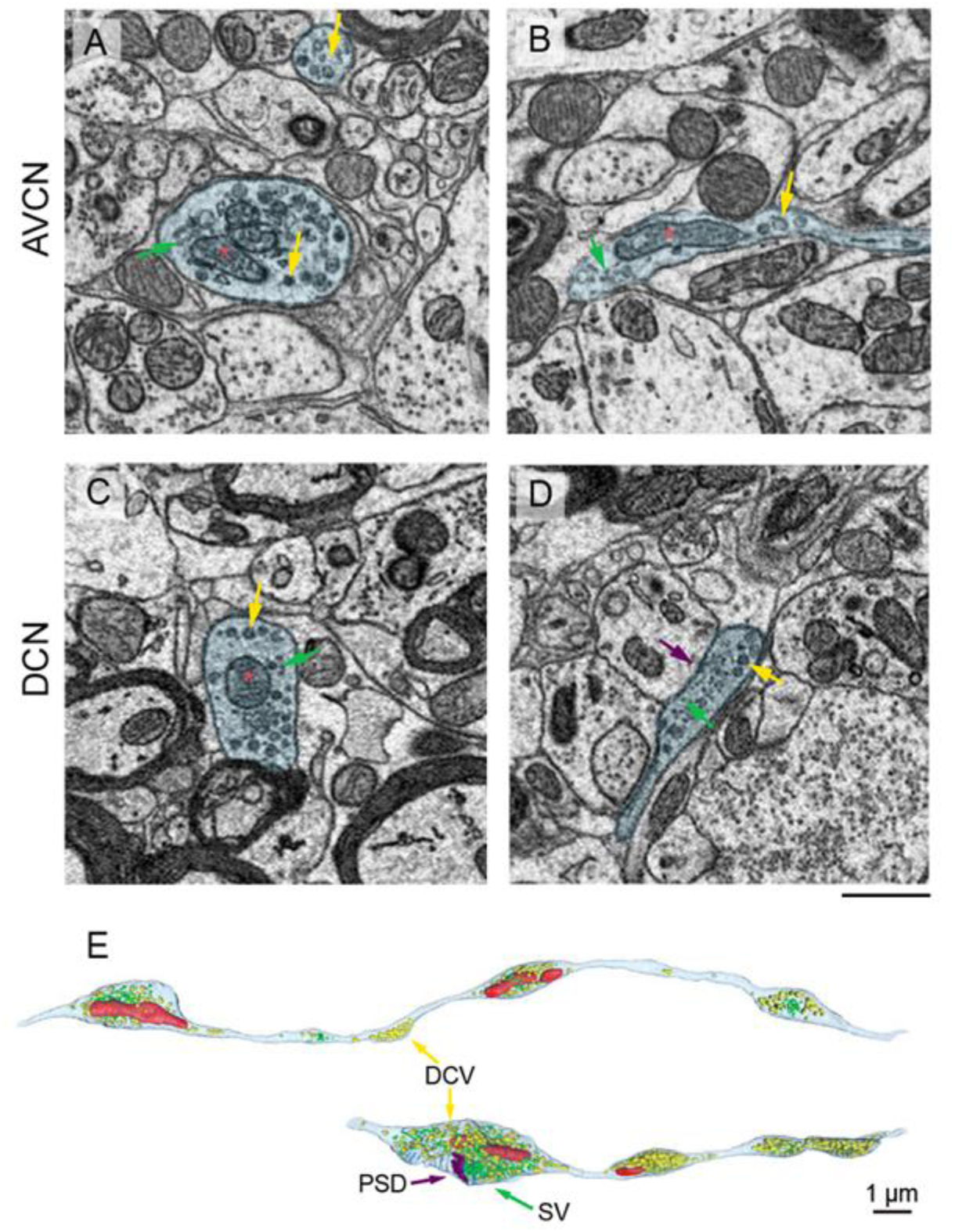
Electron microscopical evidence for putative monoaminergic varicosities in the CN. **A-D**, instances of varicosities (blue) filled with mitochondria (red asterisks), a mixture of synaptic vesicles (SVs, green arrows) and dense core vesicles (DCVs, yellow arrows). Scale bar, 500 nm, **E**, 3D reconstruction of the varicosity-forming projections (outlined with cyan) filled with DCV (yellow arrows) and synaptic vesicles (green), mitochondria (red), and putative postsynaptic density (PSD, purple). Scale bar, 1 µm.

### 3.2 Immunohistochemical verification of monoaminergic innervation in the cochlear nucleus

In order to obtain molecular evidence for monoaminergic innervation of the AVCN, we probed for the presence of the NE and 5-HT transporters, NET and SERT respectively, using STED nanoscopy. Endbulbs were labeled by staining for the vesicular glutamate transporter 1 (vGlut1, 1:2000 dilution). We observed NET positive varicosities apposed around endbulbs (Fig. 2A, 1:200 dilution, representative for n = 3 animals). We observed the SERT (Fig. 2B, 1:200 dilution, representative for n = 3 animals) in the vicinity of the endbulbs. In both cases, we observed a grainy signal forming string-like structures, smaller than 1 µm in transverse diameter, in agreement with the electron microscopical observations reported above. To verify the NET and SERT signal, we labelled slices of the locus coeruleus and the medial raphe nuclei respectively as positive controls (Fig. S1).

**Figure 2:**
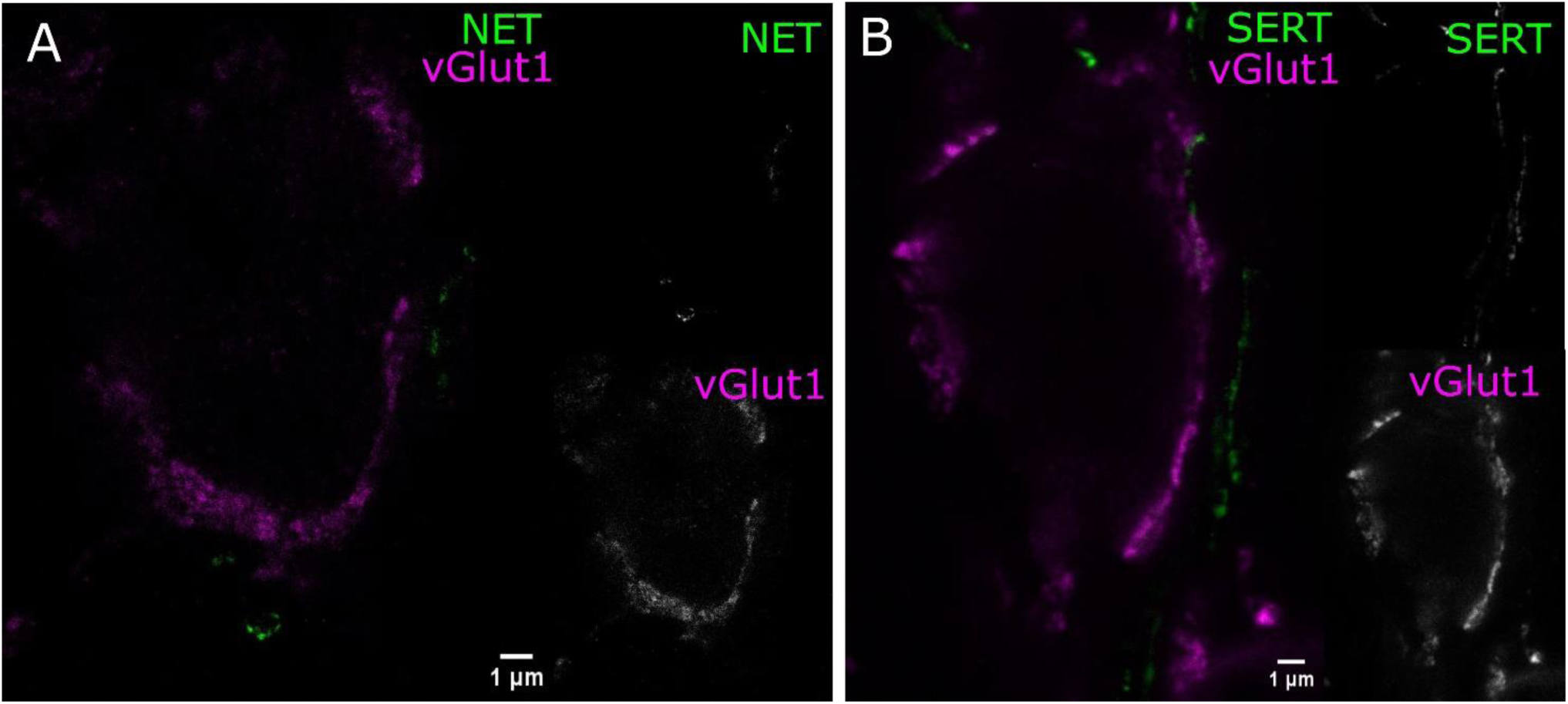
Noradrenergic and serotonergic innervation located near endbulbs of Held. Coronal sections of AVCN were used for STED imaging, A, endbulbs labelled by vGlut1 (magenta) and NET-positive structures (green), similar results from 3 animals. B, endbulbs labelled by vGlut1 (magenta) and SERT neurites (green) neighboring the endbulbs, similar results from 3 animals. The both transporter Ab display grainy signal forming strings, with transverse diameter less than 1 µm. Scale bars 1 µm.

Next, we probed for the presence of monoamine receptors in endbulbs and BCs, using vGlut1 as a presynaptic marker and Homer1 or gephyrin as excitatory or inhibitory postsynaptic markers. We shortlisted 5-HT and NE receptors according to three transcriptomic databases for SGNs (Shrestha et al., 2018; Sun et al., 2018; Li et al., 2020). Initially, we probed for β_1_-AR, β_2_-AR, β_3_-AR, α_1B_-AR, 5-HT_1A_R, 5-HT_2A_R, 5-HT_2B_R and HT_5B_. In each case, we optimized the staining protocol to improve labeling (Supplementary methods, Fig. S2). Despite the optimization, we could not detect convincing immunofluorescence except for 5-HT_5B_R (Fig. S2). In some cases, we experienced bleed-through (examples in Fig. S2). Next, we probed for the receptors 5-HT_7_R, 5-HT_2B_R, α_1B_-AR, α_1D_-AR and α_2C_-AR, again involving optimization, with a protocol that stained cryosections of immersion-fixed brainstems with the antibody against the receptor of interest (1:500) and vGlut1 (1:2000) as a presynaptic marker and Homer1 or gephyrin (1:500) as postsynaptic markers (Fig. S2). As a result, we observed a convincing staining for 5-HT_7_R and α_2C_-AR.

The α_2C_-AR immunofluorescence was mainly situated within the BC soma and their plasma membrane, possibly indicating cytoplasmic reserves or intermediate stages of receptor turnover (Fig. 3A). Less frequently, the signal was located in the vicinity of the endbulb (Fig. 3I-K). Notably, the negative control (Fig. 3E) suggested specificity of the immunolabeling. Likewise, the majority of the specific 5-HT_7_R immunofluorescence was located within the BC soma as well as in their plasma membrane (Fig. 3F, G). Occasionally, we found 5-HT_7_R signal close to the endbulbs (on Fig 3I-K). These images are close-ups on the endbulbs from a top view of the BCs. Furthermore, signal was also detected in the vicinity of the BC, possibly reflecting labelling of other neuronal structures, such as potential projections from stellate cells. The negative control (Fig. 3H) indicates specificity of the 5-HT_7_ receptor immunolabeling on Fig. 3F and 3G. Similarly, we found the receptors 5-HT_1B_ and 5-HT_4_ expressed within the BC soma (Fig. 3N and 3L respectively). The staining was performed in the same fashion except that we omitted the labelling for gephyrin which tended to off-target label structures that mostly likely represented small blood vessels. The stainings against the 5-HT_1B_R also showed signal, outlining smaller structures in proximity to, yet distinct from BCs (Fig. 3P). The 5-HT_4_R signal appeared mainly cytoplasmic. We performed negative controls, confirming the specificity of the signal (Fig. 3O and 3M, respectively). In summary, we provided positive monoamine transporter and receptor stainings, suggesting noradrenergic and serotonergic innervation of the AVCN.

**Figure 3.**
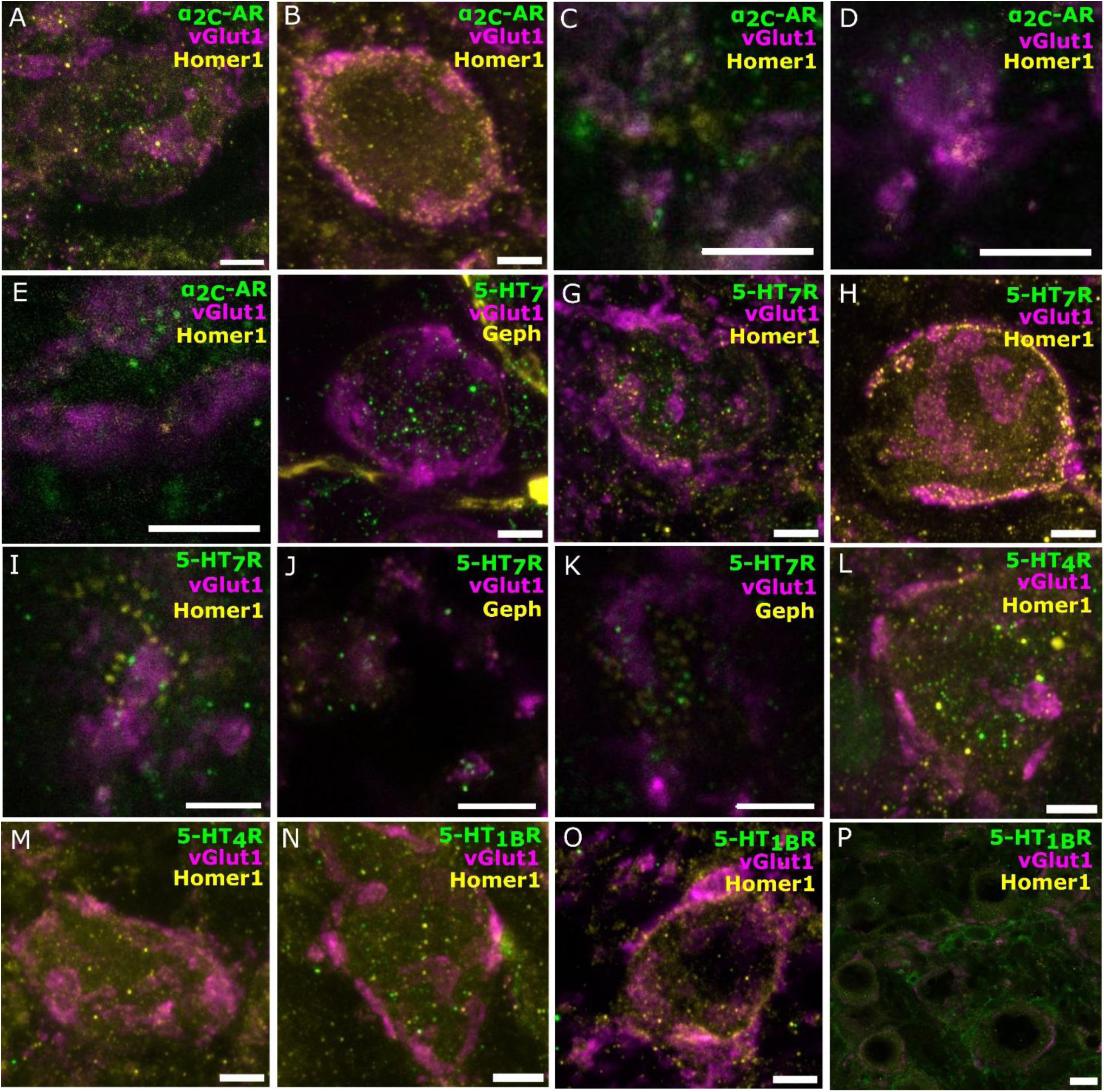
Immunofluorescence for norepinephrine and serotonin receptors in bushy cells and near endbulbs. Confocal sections of the AVCN with vGlut1 is marking the presynaptic endbulbs of Held and Homer1 or gephyrin as postsynaptic markers. **A, B**: α_2C_-AR (green, Alexa-fluor-488), vGlut1 (magenta, Alexa-fluor-647/633) and Homer1 (yellow, Alexa-fluor-568). **A**, similar results from 4 animals. **B,** a blocking peptide for the Ab against α_2C_-AR was added as a negative control. **C, D, E:** zoom-ins of the top view of BC where α_2C_-AR (green) is located closely to the endbulbs (magenta), scale bar 5 µm. **F**, 5-HT_7_R (green, Alexa-fluor-488), markers vGlut1 (magenta, Alexa-fluor-633) and Gephyrin (yellow, Alexa-fluor-568). **G, H,** 5-HT_7_ receptor (green, Alexa-fluor-488), markers vGlut1 (magenta, Alexa-fluor-647/633) and Homer1 (yellow, Alexa-fluor-568). **G.** similar results from 4 animals. **H,** is a negative control, a blocking peptide for the Ab against HT_7_R was added. **I, J, K:** zoom-ins of the top view of BC where 5-HT_7_R signal (green) is located closely to the endbulbs (magenta), scale bar 5 µm**. L, M:** 5-HT_4_ receptor (green, Alexa-fluor-488), markers vGlut1 (magenta, Alexa-fluor-647) and Homer1 (yellow, Alexa-fluor-568). **L**, similar results from 3 animals. **M**, a blocking peptide for the Ab against 5-HT_4_R was added as a negative control. **N, O, P**: 5-HT_1B_ receptor (green, Alexa-fluor-488), markers vGlut1 (magenta, Alexa-fluor-647) and Homer1 (yellow, Alexa-fluor-568) similar results from 3 animals. **O,** a blocking peptide for the Ab against 5-HT_1B_R was added as a negative control. **P**, the 5-HT_1B_R signal is also found in the vicinity of the BCs on smaller structures. Scale bars A-O 5µM. P, 10 µM.

### 3.3 Studying effects of NE or 5-HT on neurotransmission at the endbulb synapse

To elucidate the physiological basis of monoamine modulation in the AVCN, we performed whole-cell voltage-clamp recordings to capture mEPSCs from BCs in brainstem slices of C57BL/6 wild type mice in 2 mM Ca^2+^ supplemented aCSF during the third postnatal week as described in the Materials and Methods section. The mEPSCs recorded in the presence of exogenously added NE (100 µM) or 5-HT (10 µM) were compared to control data (Fig. 4). We obtained recordings from 20 cells exposed to NE for 5 minutes (Fig. 4 A, B). Our control dataset consisted of mEPSC from 21 cells in the absence of exogenously added monoamines. We observed a significantly increased frequency of spontaneous release (Fig. 4 B) in response to NE exposure (7.85 ± 0.99 Hz, n = 8538 mEPSCs, N = 20 cells from 9 mice, *p = 0.0489*) compared to control recordings (5.37 ± 0.88 Hz, n = 5414 mEPSCs, N = 21 cells from 14 mice). In a similar manner, we recorded from 16 cells (7 mice) that were incubated with a 5-HT containing solution for 5 minutes (Fig. 4 A, C). Bath application of 5-HT did not result in significantly different mEPSC frequency compared to the control (5.02 ± 0.77 Hz, n = 4821 mEPSCs, *p = 0.9511).* The amplitude and kinetic parameters of the eEPSCs remained unaltered in both cases of exogenously applied neuromodulators (data not shown).

**Figure 4.**
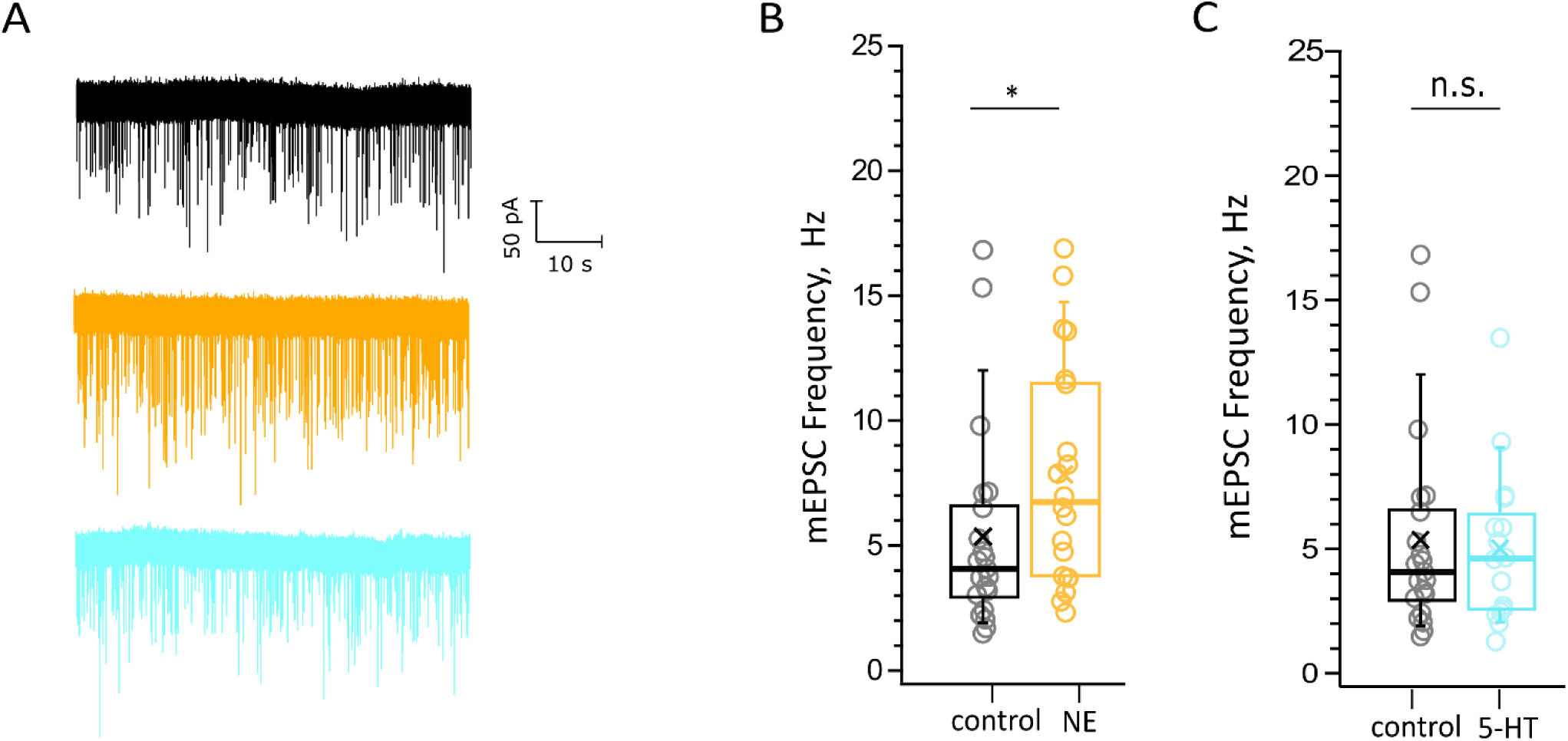
Probing for potential neuromodulation of spontaneous synaptic transmission. **A**, examples of mEPSC recordings: control (black), NE (orange), 5-HT (cyan). Scale bar 50 pA, 10s. The datasets consist of pooled recordings comparing mEPSCs of BC in a control bathing solution with mEPSCs of BC in 100 µM NE (**B**) or 10 µM 5-HT (**C**) containing solution. **B**, the mEPSC frequency is increased upon the application of NE (n = 8538 mEPSCs, N = 20 cells from 9 mice, p=0.0489). **C**, the mEPSC frequency remained unaltered upon the application of 5-HT (n = 4821 mEPSCs, N = 16 cells from 7 mice p=0,9511). The data is represented as box plots with minimum, first quartile, median, third quartile, maximum and a cross representing the mean. Each data point represents an average of the given parameter across the events recorded from one cell exposed to one condition.

To study the effect of NE on evoked synaptic transmission at the endbulb of Held synapse, we used a monopolar electrode to elicit trains of 35 monosynaptic EPSCs (eEPSCs) in BCs of p15-19 mice by extracellular stimulation of individual endbulbs. We used two stimulation frequencies, 100 Hz, for which we obtained recordings from 18 BCs from 13 mice of which a subset of 16 BCs of 11 mice also provided EPSC train data in response to stimuli delivered at 200 Hz (Fig. 5A). We observed a continuous decline in the amplitude of the eEPSCs at the different timepoints of recording under control conditions which we attribute to rundown of synaptic transmission. This can be appreciated from the difference in the amplitudes of the first eEPSCs (eEPSC_1_) of 100 Hz trains at the beginning and the end of the control condition (Fig. 5C, middle) showing a reduction to 59.3% of the initial eEPSC_1_ amplitude. Furthermore, we compared the standard deviations (SD) of the eEPSC1 amplitude for each 3 repeats in the beginning and end of the control recording and they were significantly different (SD_beginning_: 0.26 nA, SD_end_: 0.13 nA, p = 0.0026). However, the SD between the amplitudes at the end of the control recordings and NE recordings was comparable (SD_NE_: 0.15 nA, p = 0.66). Because of this rundown, we compared the last 3 recordings before and first 3 recordings after applying 100 µM NE (table 3, Fig. 5C right). The amplitude and kinetics of the eEPSC_1_ appeared unaltered upon NE application. The synaptic delay tended to be decreased without reaching statistical significance. Next, we investigated the properties of the 100 Hz and 200 Hz trains. For both frequencies the kinetics of the stereotypic endbulb short-term depression reported by the time constant (Tau (τ), Table 4) of EPSC amplitude decay during train stimulation appeared unaltered by NE. The PPR both at 100 Hz and 200 Hz in the control and NE groups were comparable and slightly facilitating in the beginning of the train. Because of the slight facilitation, we used the Elmqvist & Quastel (EQ) method (Fig. 5B) (Elmqvist and Quastel, 1965) to estimate the readily releasable pool (RRP) and the vesicular release probability (P_vr_). We did not observe differences in p_vr_ between the control and NE datasets (Table 4). The RRP of the NE dataset tended to be smaller than in control (likely reflecting the described rundown) without reaching significance (Table 4). Using a subset of our data, we compared the above-mentioned parameters for 100 and 200 Hz recordings right before and after NE application (n = 6), in order to account for the rundown and observed no significant shifts (Table S1). Additionally, we wanted to explore neuromodulation in the context of higher variability of endbulb transmission. For this purpose, we lowered the Ca^2+^ concentration to 1.3 mM. However, we did not uncover any changes in the parameters of EPSCs at 100 Hz (n = 4 cells) and 200 Hz (n = 3 cells) (Table S2). In summary, we observed an increase on the frequency of spontaneous release upon NE application, but no effect on evoked release was detected.

**Figure 5.**
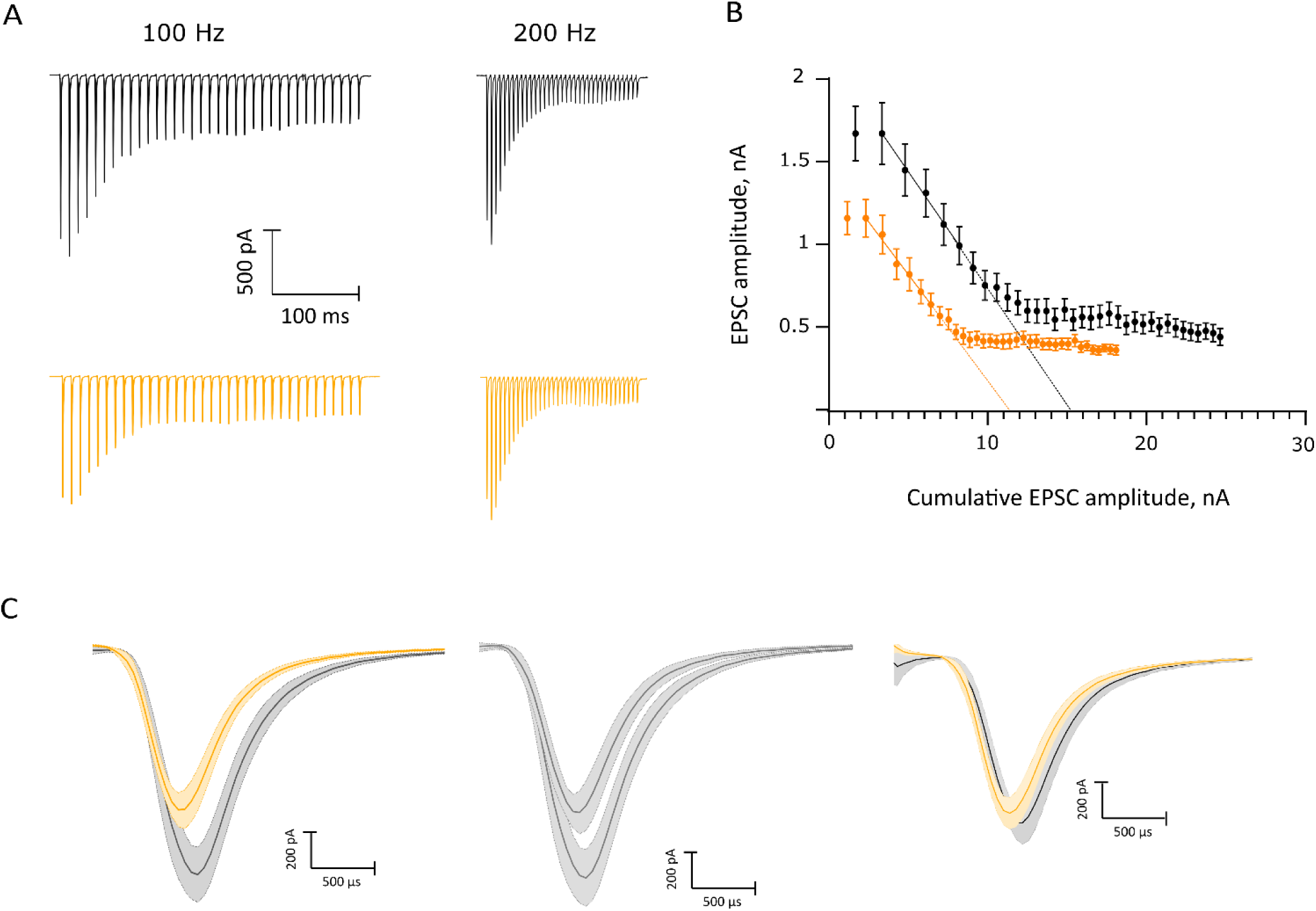
Probing for potential neuromodulation of evoked synaptic transmission. **A**, average of 100 Hz and 200 Hz trains, control recordings in black, NE recordings in orange. **B**, Elmqvist-Quastel plot of the 100 Hz NE and control trains, difference did not reach statistical significance. **C**, Rundown over time, mean values of the first EPSC amplitudes (n = 3): left, comparison of the NE and control EPSC recordings of the 100 Hz trains used for the EQ plot in (B); center, comparison of the control recordings in the beginning and end of the sampling of the control condition with a 59.3% difference; right, comparison between the recording following NE application and the control recording right before NE application. Scale bar 200 pA, 500 µs.

**Table 3.** Comparison of the first EPSC parameters before and after 100 µM NE application.

| 1st EPSC parameter (18 cells) | control | NE | p value |
| --- | --- | --- | --- |
| Amplitude, nA | 1.19±0.11 | 1.14±0.11 | 0.79 |
| Synaptic delay, ms | 1.09±0.05 | 0.97±0.05 | 0.09 |
| Rise time, ms | 0.20±0.01 | 0.22±0.03 | 0.57 |
| Decay time, ms | 0.40±0.02 | 0.37±0.02 | 0.39 |

**Table 4.**
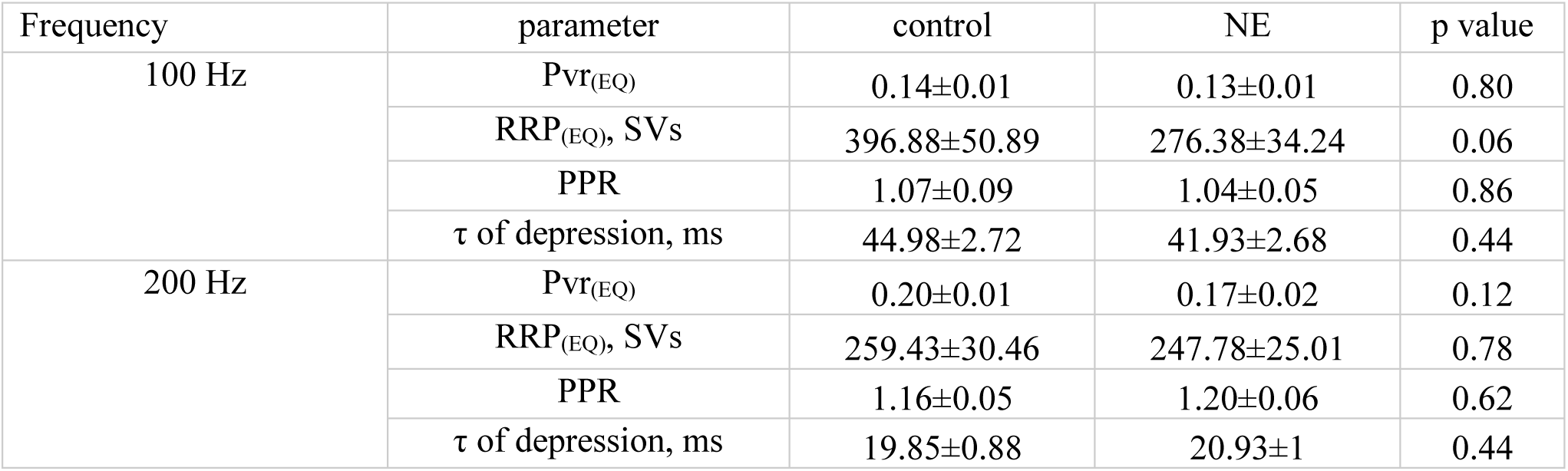
Comparison of parameters of trains of EPSCs at 100Hz and 200 Hz from the control and NE datasets.

### 3.4 Testing for noradrenergic or serotonergic modulation of BC excitability

We next probed for potential effects of NE and 5-HT on the excitability of the BC. BCs are sensitive to the speed of depolarization due to their g_KL_ and we investigated whether neuromodulation could tune this sensitivity. For this purpose, we subjected BC to currents with different rates of depolarization in current clamp experiments (Fig. 6 A_I_). We documented the slowest rate at which an AP is generated and compared it between experimental groups. Our collected datasets consisted of recordings from 19 cells from 14 mice in the control condition, 14 cells from 7 mice in the 5-HT group (10 µM) and 17 cells from 7 mice exposed to NE (100 µM). The average rate of depolarization in the NE dataset was 2.60 ± 0.24mV/ms (Fig. 6 A_II_), showing a non-significant trend of firing APs at slower rates of depolarization compared to the control recordings (*p = 0.2913*). The average rate of depolarization appeared to be quite similar between the control (2.97 ± 0.24mV/ms) and 5-HT groups (2.99 ± 0.27 mV/ms *p = 0.9284,* Fig. 6 A_III_).

**Figure 6.**
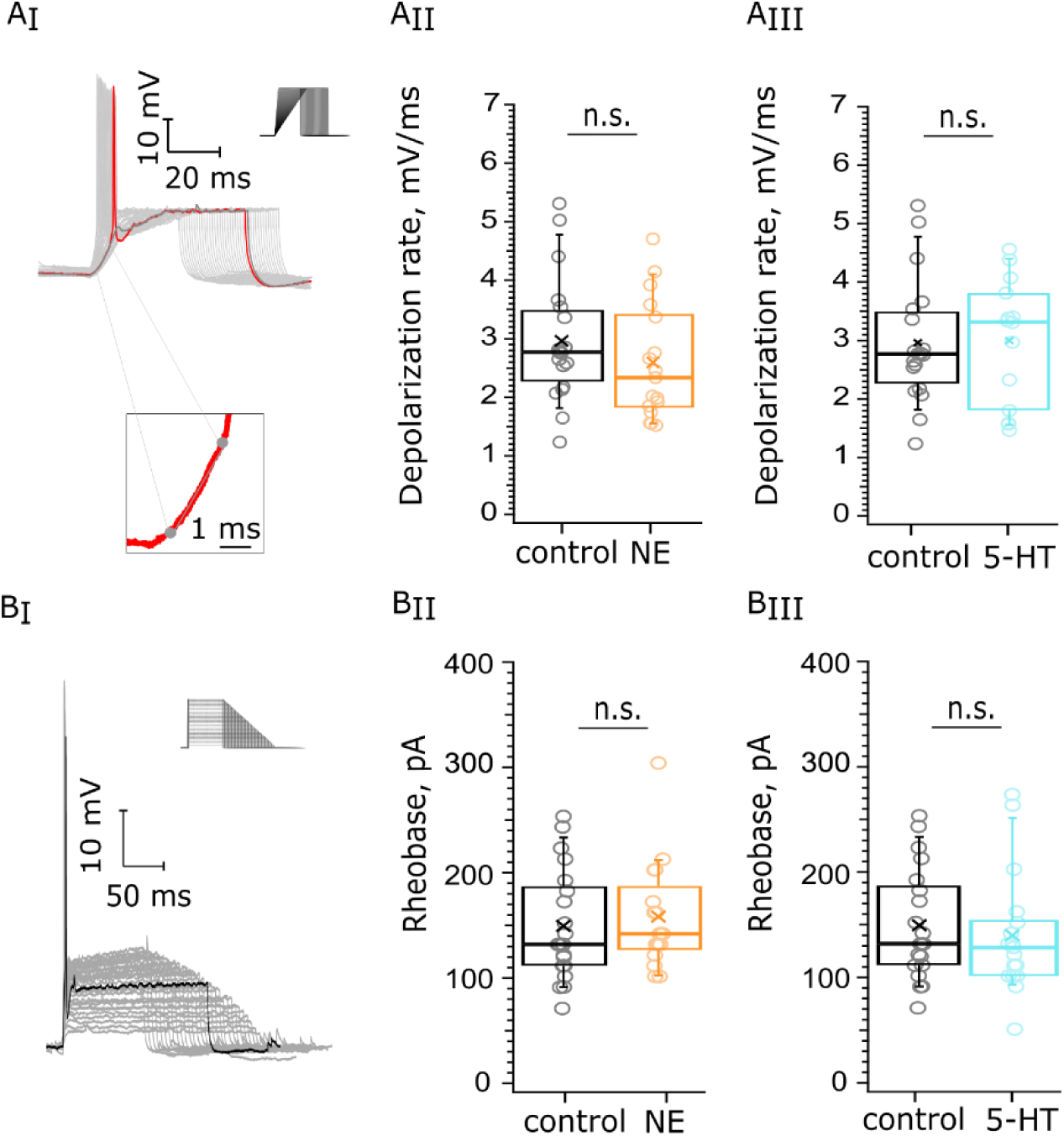
Testing for neuromodulation of BC excitability. 10 µM 5-HT or 100 µM NE were applied for 5 min following recordings under control conditions: **A**, BCs’ rate of depolarization, 19 cells (14 mice) in the control dataset, 14 cells (7 mice) for 5-HT and 17 cells (7 mice) for NE. **A_I_**, we documented the slowest rate at which an AP is generated (red trace) and estimated the derivative of the linear component of the voltage trace before the generation of the AP. Scale bar 10 mV, 20 ms. **A_II_**, the rate of depolarization was tended to be lower upon NE application without reaching significance (p = 0.2913). **A_III_**, the rate of depolarization was not significantly altered upon 5-HT administration (p = 0.9284). **B**, Rheobase of BCs. Our datasets consisted of recordings from 20 cells, 13 mice in the control dataset, 17 cells, 7 mice for 5-HT one and 16 cells, 7 mice for NE. Scale bar 10 mV, 50 ms **B_I_**, we injected currents with different amplitudes and duration (top right corner) and documented the first current value at which an AP (black) was generated. **B_II_**, the Rheobase did not display significant change upon NE application (p = 0.5772 or **B_III_**, upon 5-HT administration, p = 0.4734)

Next, we aimed to examine whether the excitability of the BCs was altered upon NE or 5-HT application. For this purpose, we quantified the rheobase - the minimal current amplitude of infinite duration that leads to the crossing of the depolarization threshold of the cell membrane (Fig. 6 B). We obtained a dataset of control recordings from 20 cells among 13 mice as well as recordings from cells, exposed for 5 minutes to 5-HT (17 cells, 7 mice) and to NE (16 cells, 7 mice). The average Rheobase of the control recordings was 149.52±11.54 pA. Similarly, the rheobase of the NE recordings (158.4 ± 12.7 pA, p = 0.5772, Fig. 6 B_II_) and the Rheobase of the 5-HT recordings (139.9 ± 13.8 pA, *p = 0.4734*, Fig. 6 B_III_) showed similar values in comparison with the control. In conclusion, neither 100 µM NE nor 10 µM 5-HT affected depolarization speed and rheobase of BCs.

Additionally, we delivered trains of action potentials by postsynaptic stimulation of the BCs at 100 Hz and 200 Hz. We did not observe significant shifts in the first AP kinetics, the spike probability at 200 Hz or the amplitude of the APs over the course of the 100 Hz trains for both 5-HT and NE (Fig. S3-S6).

### 3.5 Probing for noradrenergic or serotonergic modulation of outward currents of BCs

Next, we probed for potential effects of NE and 5-HT on the currents mediated by low-voltage activated K^+^ channels (I_KL_, reference, Materials and Methods). We applied depolarizing voltage steps (5 mV) from −90 mV to −40 mV activating voltage-sensitive outward current and added blockers of to isolate K^+^ currents (10 µM ZD7288, 1 µM TTX, 0,25 mM CdCl2, 10 µM DNQX, 2 µM strychnine and 10 µM bicuculline). The peak of the I_KL_ positive currents has been shown to be at −40 mV (Fu et al., 2021), so we focused on currents recorded at −40 mV (Fig. 7A top, darker trace) for our analysis. To estimate what portion of this peak outward current densities at −40 mV is attributed to I_KL_ we applied α-dendrotoxin (α-DTX, 50 nM) which blocks Kv_1.1_, Kv_1.2_ and Kv_1.6_ channels. This solution was perfused for 9 minutes following control recordings. We observed a 44.87% reduction of the outward current densities at −40 mV (Fig. 7A bottom). We then probed for potential effects of application of 100µM NE and 10 µM 5-HT, respectively. We did not detect significant differences upon application of either neuromodulator. The average outward current density (101.6 ± 5.7 pA/pF from 18 cells, 7 mice, *p = 0.2017,* Fig. 7 B) at 5 minutes of 100µM NE perfusion tended to be increased compared to the density of the control dataset (91.0 ± 4.4 pA/pF from 17 cells, 8 mice) without reaching statistical significance. The average current density of the control and 5-HT datasets was comparable: 5-HT dataset: 95.9 ± 8.5 pA/pF from 14 cells, 7 mice, *p = 0.6791,* Fig. 7 C). In conclusion, we did not observe changes in the I_KL_ peak current density while pharmacologically applying 10 µM 5-HT or 100 µM NE.

**Figure 7.**
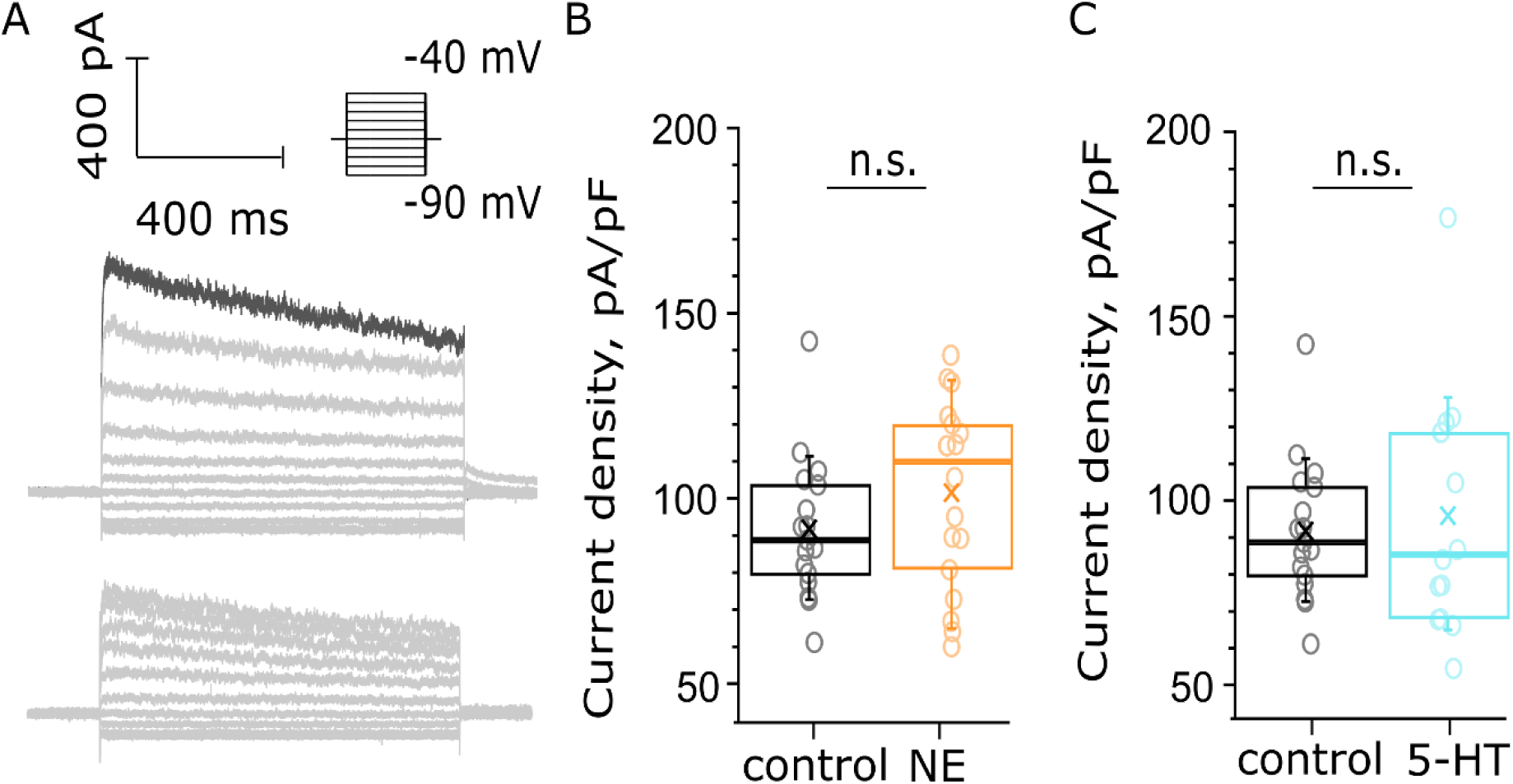
Probing for noradrenergic or serotonergic modulation of outward currents of BCs. **A**, an example recording of positive outward current in BC. Scale bar 400 pA, 400 ms. Depolarizing voltage steps (5mV) from −90 mV to −40 mV activated voltage-sensitive outward current. Top: Recordings were made in the presence of 10 µM ZD7288, 1 µM TTX, 0,25 mM CdCl_2_, 10 µM DNQX, 2 µM strychnine and 10 µM bicuculline. Bottom: a negative control for I_KL_, addition of 50 nM α-DTX to the solution reduced the outward current at −40 mV with 44.87%. **B, C** the current density was calculated as the ratio between the measured current at −40 mV of each recording and the C-slow capacitance of each cell. **B**, the peak current density was non-significantly increased by NE perfusion (*p = 0.2017*, n_control_ = 17 cells from 8 mice; n_NE_ = 18 cells from 7 mice). **B**, the peak current density was unaltered by 5-HT perfusion (*p=0.6791*, n_control_ = 17 cells from 8 mice; n_5-HT_ = 14 cells, 7 mice).

### 3.6 Probing for noradrenergic or serotonergic modulation of hyperpolarization activated currents of BCs

Finally, given the immunohistochemical indication of the presence of α_2C_-AR, 5-HT_1B_R, 5-HT_4_R and 5-HT_7_R in BCs which modulate cAMP levels, we decided to probe for effects of NE or 5-HT on hyperpolarization induced (I_h_) currents that are modulated cAMP. Voltage steps (5mV) from −112 mV to −57 mV activated voltage-sensitive inward current at the hyperpolarizing voltages in the presence of 25 nM α-DTX, 1 µM TTX, 0,25 mM CdCl2, 10 µM DNQX, 2 µM strychnine and 10 µM bicuculline (Fig. 8 A). Our recorded traces were sorted in the following groups: a control group, an NE group, comprising recordings from cells exposed to 100 μM NE for 5 minutes and finally a 5-HT group, subjected to 5 minutes of bath-applied 10μM of 5-HT. To estimate what portion of this negative current is attributed to I_h_ we performed a negative control. First, we recorded the outward current using our control solution, which was followed by perfusion of a solution, supplemented with 10 µM ZD7288 to block HCN channels. This second solution was perfused for 9 minutes and the difference in currents can be appreciated on Fig. 8A. We observed a 48.49% reduction of the inward current at −112 mV, which could be attributed to the blocking of HCN channels. The average inward current density estimated as the plateau at −112 mV of control dataset recordings (−35.52 ±4.04 pA/pF from 13 cells, 7 mice) was comparable to that yielded from both the 5-HT (−38.69±3.76 pA/pF from 14 cells, 6 mice, *p = 0.5850*, Fig. 8C) and NE (−36.87 ±4.83 pA/pF from 12 cells, 5 mice, *p = 0.6280*, Fig. 8B) datasets. To conclude, we did not detect changes in I_h_ upon administration of 10 µM 5-HT or 100 µM NE.

**Figure 8.**
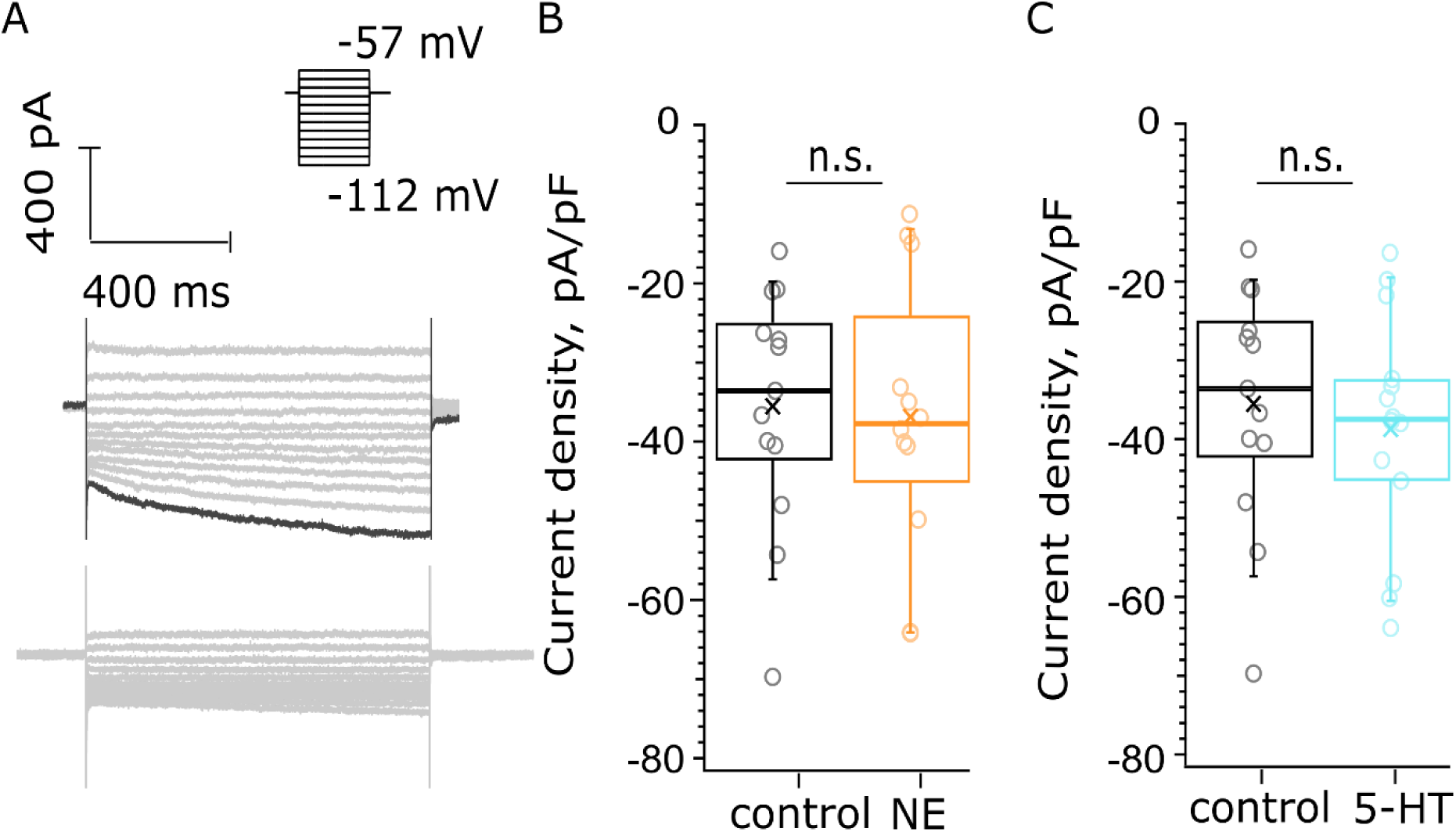
Probing for noradrenergic or serotonergic modulation of hyperpolarization activated currents of BCs. **A**, voltage steps (5mV) from −112 mV to −57 mV activated voltage-sensitive inward current at the hyperpolarizing voltages. Scale bar 400 pA, 400 ms. Top: The recordings were made in the presence of 25 nM α-DTX, 1 µM TTX, 0,25 mM CdCl_2_, 10 µM DNQX, 2 µM strychnine and 10 µM bicuculline. Bottom: negative control for I_h_ currents, addition of 10 µM ZD7288 reduced the inward currents with 48.49%. The amplitude of the inward current that we measured was derived as the average current amplitude at the last 100 ms of the voltage trace recorded at −112 mV. **B, C,** the current density was calculated as the ratio between the measured current at −112 mV of each recording and the C-slow capacitance of each cell. **B**, the current density was unaltered in the presence of 100 µM NE after 5 min of perfusion (*p=0.6280*, n_control_ = 13 cells from 7 mice; n_NE_ = 12 cells from 5 mice). **C**, we did not observe a significant difference in the current density in the presence of 10 µM 5-HT at 5 mins of perfusion (*p = 0.5850*, n_control_=13 cells from 7 mice; n_5-HT_ = 14 cells, 6 mice).

## 4 Discussion

In this study we provide initial morphological and functional evidence for monoaminergic innervation of the CN. We found varicose neurites featuring dense core and synaptic vesicles in volume EM reconstructions made for the AVCN and DCN which are compatible with monoaminergic innervation. Moreover, we obtain immunohistochemical proof of monoamine transporters being present in the CN and explored the receptor complement of such neurotransmitters. Our report of expression of a_2C_-AR as well as 5-HT_7_, 5-HT_1B_, and 5-HT_4_ receptors in the AVCN provide support for volume neuromodulatory transmission affecting this early auditory circuit. Finally, we found a subtle increase in the mEPSC frequency of BCs upon application of 100 µM NE suggesting a direct noradrenergic modulation of the endbulb of Held. Yet, much remains to be done to rigorously analyze the functional role of monoaminergic innervation of the CN beyond this first preliminary study.

### 4.1 Morphological evidence for the presence of varicosity-containing projections in the cochlear nucleus

Here we provided immunohistochemical evidence for the presence of the NET and SERT in the vicinity of the endbulb-BC synapse. To further consolidate this finding of monoamine releasing varicosities we turned to SBEM and found varicosity-like neurites in the DCN and the AVCN. Indeed, a previous electrophysiological study in the mouse DCN demonstrated serotonergic innervation acting through 5-HT_2_ and 5-HT_7_ receptors (Tang and Trussell, 2015). Further studies should extend the volume EM analysis to larger volumes and also employ immuno-EM experiments to test the correspondence of these varicosities to 5-HT neurons. Our preliminary investigation of monoamine receptors in the AVCN revealed initial evidence of 5-HT_1B_, 5-HT_4_, 5-HT_7_ receptors and the α_2C_-adrenergic receptors inside BCs and near their plasma membrane. The intracellular localization of the receptors could be explained by GPCR internalization as a result of stimulation. This mechanism of action has been described for the 5-HT_2A_R, where after stimulation, the GPCR colocalizes with endosome markers such as Rab5 and Rab7 (Eickelbeck et al., 2019). Additionally, we observed 5-HT_7_R and α_2C_-AR signal in the vicinity of the endbulbs in some of the BCs that we imaged. However, the use of higher resolution imaging, such as STED microscopy, will be required to further and more accurately confine the localization of the receptors relative to pre- or postsynaptic elements.

The 5-HT_7_R is coupled to G_s_ proteins and its activation raises cAMP levels which in turn trigger PKA – dependent effector cascades. The 5-HT_7_ receptor is robustly expressed in the thalamus and hypothalamus, as well as the hippocampus and cortex. This receptor is involved in thermoregulation, circadian rhythm, learning and memory, sleep and mood regulation. Similarly, the 5-HT_4_R is a G_s_ coupled receptor that is involved in learning, memory and mood regulation and is expressed in the cortex and hippocampus, as well as limbic regions, namely in the basal ganglia and amygdala. On the contrary the 5-HT_1B_R is coupled to G_i_ proteins and expressed in the cortex as well as basal ganglia. We observed the 5-HT_1B_R signal in our stainings in structures localizing near BC. This suggests that other cell types are regulated by 5-HT signaling in the AVCN. Yet, complimentary electrophysiological experiments will be required to elucidate the mechanism of action of 5-HT_7_, 5-HT_4_ and 5-HT_1B_ receptors in BC.

α_2C_-ARs are known to serve both as autoreceptors, controlling noradrenaline release, and as heteroreceptors. α_2C_-AR heteroreceptors were shown to regulate 5-HT transmission by inhibition of 5-HT release, but to a lesser extent than α_2A_-AR (Scheibner et al., 2001). Concentrations of NE, and 5-HT were shown to be increased in the brains of α_2C_-AR knock-out mice (Sallinen et al., 1997). In this context, the presence of the α_2C_-AR in the vicinity of the endbulb synapse may indicate an intricate regulation network of this early integration station of the auditory pathway. As for 5-HT_7_ receptors, further work is required to dissect the mechanism of action of 5-HT in the AVCN.

Additionally, other receptor subtypes might be present, but we might have failed to detect them under the chosen conditions and with the commercial antibodies at our discretion. A way to tackle this could be the use of melanopsin-variants called Camello-XR ((Eickelbeck et al., 2019)). These are G-protein-coupled opsins, carrying a fluorescent label (e.g. mCherry or eGFP). The C-termius of these proteins can be modified to contain an amino acids sequence from the C-terminal region of a neuromodulator receptor (e.g. 5-HT_2A_), which is a localization signal that leads to the trafficking of the protein to the membrane domains that are normally occupied by the known receptor.

### 4.2 Neuromodulator pharmacology and synaptic transmission in the AVCN

We observed an augmented mEPSC frequency in the presence of 100 µM NE which reflects a higher number of SV spontaneously fusing with the membrane. Potentially, this could be attributed to the priming of synaptic vesicles and transitions from lose to tight docking state (Neher and Brose, 2018), further explainable by an overall increase of available vesicles at AZs (Patzke et al., 2019), yet without affecting the kinetics of state transitions. However, in order to argue in favor of such speculations we need a mechanistic proof for such a process. Our evoked release data did not show significant shifts in the PPR, P_vr_, RRP, nor in the amplitude of the EPSCs or the τ of depression. There are indications for differential regulation of spontaneous and evoked release (Ramirez and Kavalali, 2011). Hence, there is a possibility that the increased frequency of spontaneous release upon NE application is governed by a regulation mechanism separate from the ones controlling evoked release. Additionally, we did not uncover a significant shift in the BCs’ characteristic ion conducting channels. It is possible that these channels are in fact not regulated by neuromodulators and/or that neuromodulation operates on AVCN processes that we did not analyze. A recent study revealed that cAMP reduced the amplitude of APs and the speed of their propagation over long axonal distances due to sodium current inhibition (Abate et al., 2024.). This could potentially serve as a tuning mechanism of transmission between the SGNs and BCs. However, in our preliminary AP train experiments with a dataset of 5 cells we did not observe a significant change is APs’ size over the course of the train. Still, the non-physiological pharmacological application of monoamines and/or the limited size of the datasets might have prevented detection of neuromodulatory effects.

## 5 Limitations of the present study

In order to further probe for monoaminergic modulation of the endbulb synapse, more specific stimulation or inhibition of endogenous monoaminergic modulation of the AVCN should be used. This could be achieved through optogenetic control of the putative monoamine projections in the AVCN. In future studies we aim to expand our work in this direction through AAV-mediated cell-type specific expression of excitatory and inhibitory ChRs using appropriate Cre-mice such as the Dbh-cre and SERT-cre lines. Additionally, the AVCN is a complex structure and we cannot rule out the possibility that synapses other than the one we studied could be undergoing neuromodulation. Furthermore, our immunohistochemistry data needs to be backed up with functional evidence for the action of NE and 5-HT at the endbulb – BC synapse. In our EM study we uncovered neurites that could potentially be of monoaminergic origin, however this is yet to be confirmed. Perhaps post-embedding immunogold EM or even FRIL using robust antibodies against surface molecules such as SERT or NET could elucidate the precise localization of noradrenergic neurites relative to known AVCN circuit elements.

## 6 Conflict of Interest

The authors declare that the research was conducted in the absence of any commercial or financial relationships that could be construed as a potential conflict of interest.

## 7 Author Contributions

The experimental work was performed by M.G. (acute slice electrophysiology, immunohistochemistry), T.A. (electrophysiology, immunohistochemistry) and Y.Q., F.W., Y.H. and C.W. (electron microscopy). M.G. and T.M. prepared the manuscript with contributions from all authors.

## 8 Funding

This work was supported by the Deutsche Forschungsgemeinschaft (DFG) through the collaborative research center 1286 to T.M. and C.W. and Cluster of Excellence Multiscale Bioimaging (EXC2067, MBExC) to T.M. as well as by Fondation Pour l’Audition (FPA RD-2020-10, TM). The EM experiment was funded by Innovative Research Team of High-level Local Universities in Shanghai (SHSMU-ZLCX20211700).

## 9 Acknowledgments

We thank S. Gerke, I. Herfort, and C. Senger-Freitag for expert technical assistance, P. Räke-Kügler, K. Dinter and I. Herfort for administrative help and J. Neef for help with the STED imaging and image analysis.

## Notes

### Competing Interest Statement

The authors have declared no competing interest.

