## Supplementary material for "Neuromodulation of the endbulb of Held synapse in the cochlear nucleus"

**Supplement**

Supplementary methods

Fig. S1 Positive control of NET and SERT immunolabeling in Locus coeruleus and Raphe nuclei.

Fig. S2 Immunolabeling for 5-HT and NE receptors, optimization of the staining protocol.

Table S1 Analysis of EPSCs trains evoked by 100Hz and 200 Hz of stimulation in control and NE conditions at 1.3 mM $Ca^{2+}$.

Table S2 Analysis of EPSCs trains evoked by 100Hz and 200 Hz of stimulation right before and during NE application.

Fig. S3 Probing for effects of 5-HT on the first AP at 100Hz stimulation frequency.

Fig. S4 Probing for effects of NE on the first AP at 100Hz stimulation frequency.

Fig. S5 Relative amplitude of the last AP of BC firing during 100 Hz train stimulation in control, 5-HT and NE conditions.

Fig. S6 BC spike probability at 200 Hz.

**Supplementary methods – optimization of the staining protocol**

Our survey started with a parallel probing for $\beta_3$-AR (S1, G) and 5-$HT_{2A}R$ (S1, C) using the primary antibodies against these receptors at a dilution of 1:200. The receptors in these cases were marked with a secondary Ab Alexa Fluor 568, while the markers V-Glut1(1:1000) – with Alexa Fluor 488 and Gephyrin (1:500) – with Alexa Fluor 633. We acquired z-stacks of each BC and presented them as maximum intensity projections.

What can be readily observed in these experiments is strong bleed-through from the 488 nm channel to the 568 nm one. Apart from the bleed-through overlap, according to the negative controls the signal representing the receptors of interest showed a small grainy dotted pattern localized both inside and around the cells. Additionally, the signal from the 633 nm channel was not entirely specific for Gephyrin, but also delineated blood vessels (unspecific binding).

Consequently, we performed stainings against $\beta_1$-AR (S1, E), $\beta_2$-AR (S1, F), 5-$HT_{1A}R$ (S1, B) and 5-$HT_{5B}R$ (S1, A) in dilution 1:200 and using a secondary Ab with peak emission 488 nm wavelength, the marker vGlut1 – with peak at the 568 nm wavelength and using a different secondary Ab against gephyrin, this time conjugated with Alexa Fluor 633 nm. However, our Gephyrin staining was still unsuccessful. Our staining against the 5-$HT_{5B}R$ displayed a strong signal and no bleed-though. In the stainings against the β1-AR and 5-$HT_{1A}R$, bleed-through was still present, even from the 568 nm to the 488 nm channel. The β2-AR staining showed localization of the Ab on the nuclear membrane.

We used the staining against β3-AR as a reference for further optimization. To avoid bleed-through we conjugated the anti-β3-AR Ab with Alexa Fluor 488 and vGlut1 – with Alexa Fluor 633. However, even in this configuration where the two markers were placed far from each other spectrally bleed-through was still present. We also tried to increase the anti-β3-AR Ab concentration to 1:100, due to the concern that bleed-through was more relevant for a faint signal. However, this did not improve the quality of the images but rather more unspecific binding occurred.

Therefore, we decided to decrease the concentration to 1:500 and probe for new receptors, to expand our dataset, since the studied ones so far might not display prominent or any expression in the vicinity of the EoH – BC synapse. Additionally, we decided to replace the postsynaptic marker Gephyrin with Homer1, because of the observed unspecific binding to capillaries. The new batch of receptors tested included 5-$HR_7R$ (Fig. 2 B, C), 5-$HT_{2B}R$ (S1, D), $\alpha_{1B}$-AR (S1, H), $\alpha_{1D}$-AR (S1, I) and $\alpha_{2C}$-AR (Fig. 2 A). The stainings against the 5-$HR_7R$ and $\alpha_{2C}$-AR receptors showed a high intrinsic signal and no bleed-through was observed in those probes. This led us to the conclusion that the specific immunofluorescence signal in the previous stainings displaying bleed-through (examples on Fig. S1) was too dim such that the high gain setting being used during image acquisition emphasized the background signal.

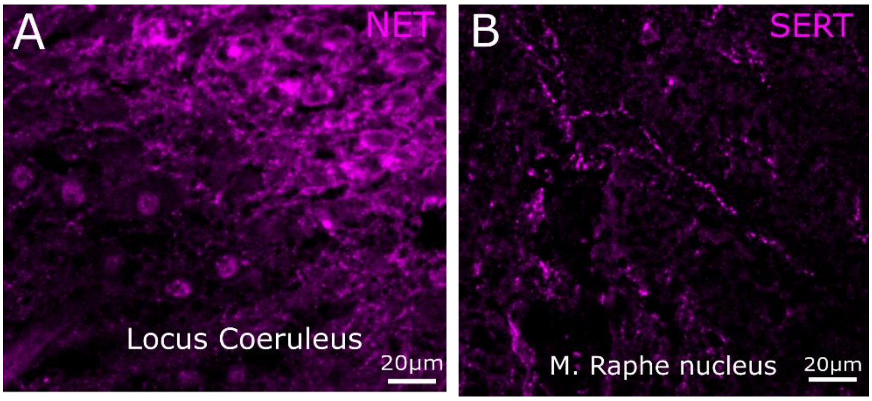


**Figure S1 Positive control of NET and SERT immunolabeling in Locus coeruleus and Raphe nuclei**: A, NET staining in Locus coeruleus. B, SERT staining in the medial Raphe nucleus.

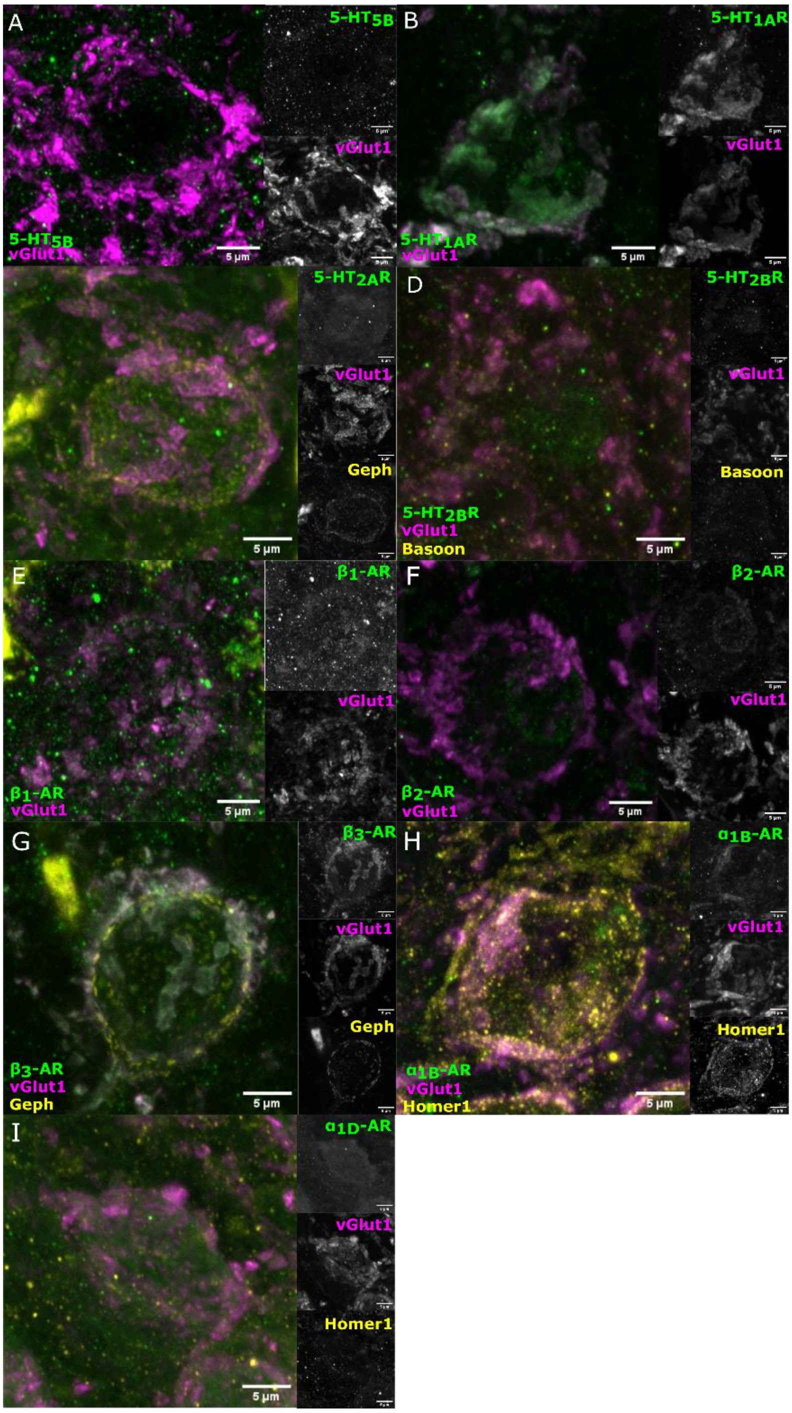


**Figure S2 Immunolabeling for 5-HT and NE receptors, optimization of the staining protocol.** All antibodies were tested with a negative control containing the binding peptide of the monoamine receptor AB (data not shown). A, 5-HT$_{5B}$ receptor (green, Alexa-fluor-488), vGlut1 (magenta, Alexa-fluor-568). B, HT$_{1A}$R (green, Alexa-fluor-488), vGlut1 (magenta, Alexa-fluor-568). C, 5-HT$_{2A}$R ( green, Alexa-fluor-488), vGlut1 (magenta, Alexa-fluor-568) and gephyrin (yellow, Alexa-fluor-633). D, 5-HT$_{2B}$R (Alexa-fluor-488), vGlut1 (magenta, Alexa-fluor-568), bassoon (yellow, Alexa-fluor-633). E, β$_1$-AR (green, Alexa-fluor-488), vGlut1 (magenta, Alexa-fluor-568). F, β$_2$-AR (Alexa-fluor-488), vGlut1 (magenta, Alexa-fluor-568). G, β$_3$-AR (green, Alexa-fluor-568), vGlut1 (magenta, Alexa-fluor-488) and gephyrin (yellow, Alexa-fluor-633). H, α$_{1B}$-AR (green, Alexa-fluor-488), vGlut1 (magenta, Alexa-fluor-568) and Homer1 (yellow, Alexa-fluor-633). I, α$_{1D}$-AR (green, Alexa-fluor-488), vGlut1 (magenta, Alexa-fluor-568) and Homer1 (yellow, Alexa-fluor-633).

**Table S1 Analysis of EPSCs trains evoked by 100Hz and 200 Hz of stimulation in control and NE conditions at 1.3 mM $Ca^{2+}$.** N=6 cells from 4 animals

| frequency | parameter | control | NE | p value |
|---|---|---|---|---|
| 100 Hz | $Pvr_{(EQ)}$ | 0.15±0.01 | 0.10±0.01 | 0.05 |
| | $RRP_{(EQ)}$, SVs | 404.28±69.65 | 340.19±75.37 | 0.09 |
| | PPR | 1.12±0.07 | 0.94±0.05 | 0.58 |
| | τ of depression, ms | 39.64±5.59 | 47.03±7.44 | 0.36 |
| 200 Hz | $Pvr_{(EQ)}$ | 0.16±0.03 | 0.12±0.02 | 0.27 |
| | $RRP_{(EQ)}$, SVs | 340.74±48.64 | 333.39±37.65 | 0.92 |
| | PPR | 1.12±0.09 | 1.10±0.08 | 0.92 |
| | τ of depression, ms | 35.04±11.09 | 25.37±3.86 | 0.44 |

**Table S2 Analysis of EPSCs trains evoked by 100Hz and 200 Hz of stimulation right before and during NE application.** N=4 cells from 4 animals for the 100 Hz recordings, n=3 cells from 3 animals for the 200 Hz recordings

| Frequency | parameter | control | NE | p value |
|---|---|---|---|---|
| 100 Hz | $Pvr_{(EQ)}$ | 0.09±0.01 | 0.10±0.02 | 0.88 |
| | $RRP_{(EQ)}$, SVs | 202.57±80.82 | 195.10±82.16 | 0.96 |
| | PPR | 1.49±0.25 | 1.19±0.11 | 0.37 |
| | τ of depression, ms | 68.04±7.72 | 77.15±21.77 | 0.75 |
| 200 Hz | $Pvr_{(EQ)}$ | 0.15±0.03 | 0.08±0.02 | 0.30 |
| | $RRP_{(EQ)}$, SVs | 175.42±67.43 | 333.47±109.85 | 0.37 |
| | PPR | 1.84±0.33 | 1.95±0.06 | 0.7 |
| | τ of depression, ms | 44.37±8.87 | 48.32±13.01 | 0.85 |

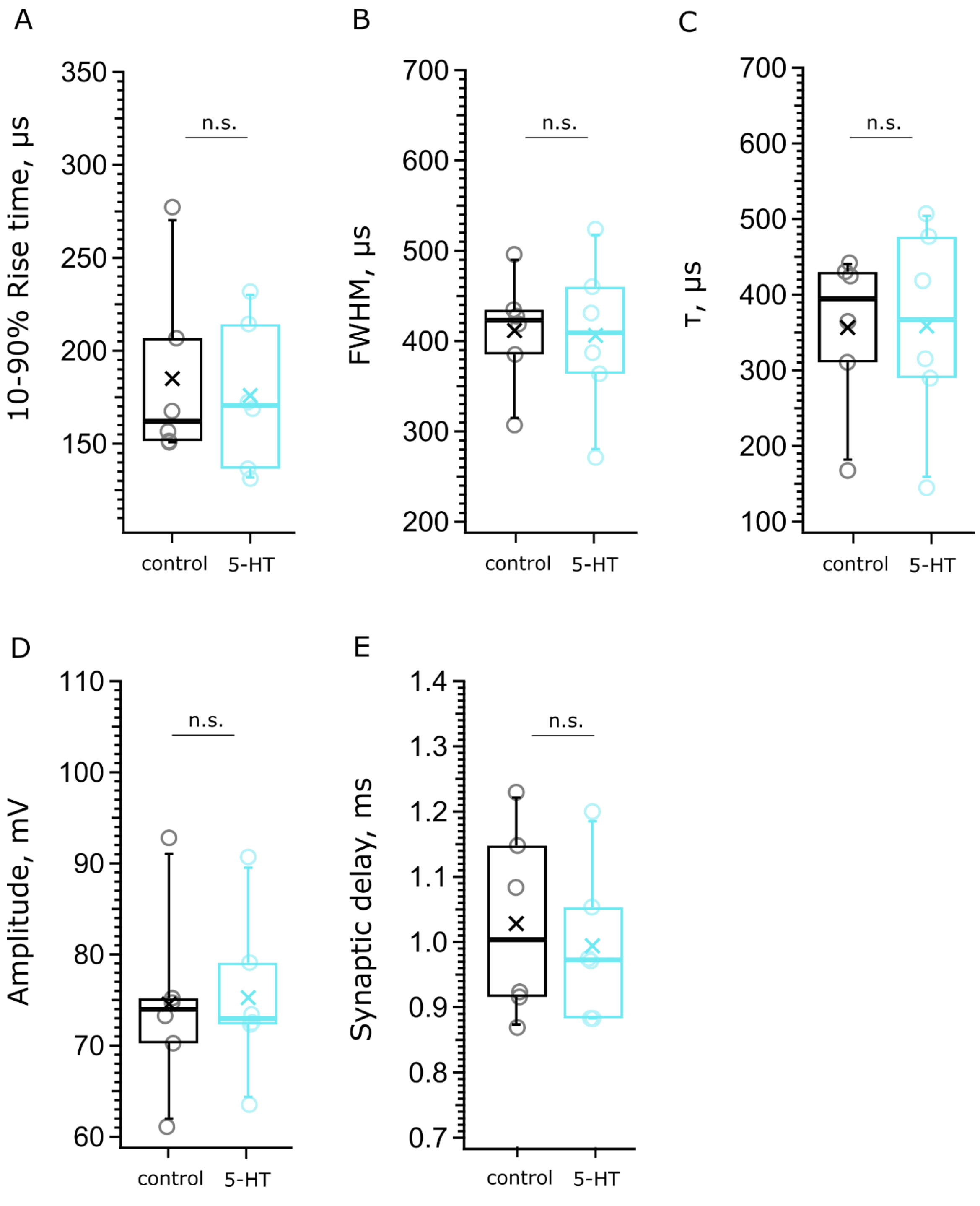


**Figure S3 Probing for effects of 5-HT on the first AP at 100Hz stimulation frequency.** We performed current-clamp recordings of BC action potentials (APs) in response to train stimulations delivered to the afferent ANFs with a monopolar electrode. We first acquired AP trains in the control solution (black), followed by recording responses to the same sequence of stimuli, while bathing the calls with the 10 µM 5-HT solution (blue). N=6 cells from 6 mice A, 10-90% rise time of the AP, *p=0. 7105*, B – FWHM of the AP, *p=0.9045*, C, τ, *p=0.4670*, D, Amplitude of the AP, *p=0.5722*, E, Synaptic delay, *p=0.8815*. Each data point represents an average of the given parameter across the events in one mEPSC recording.

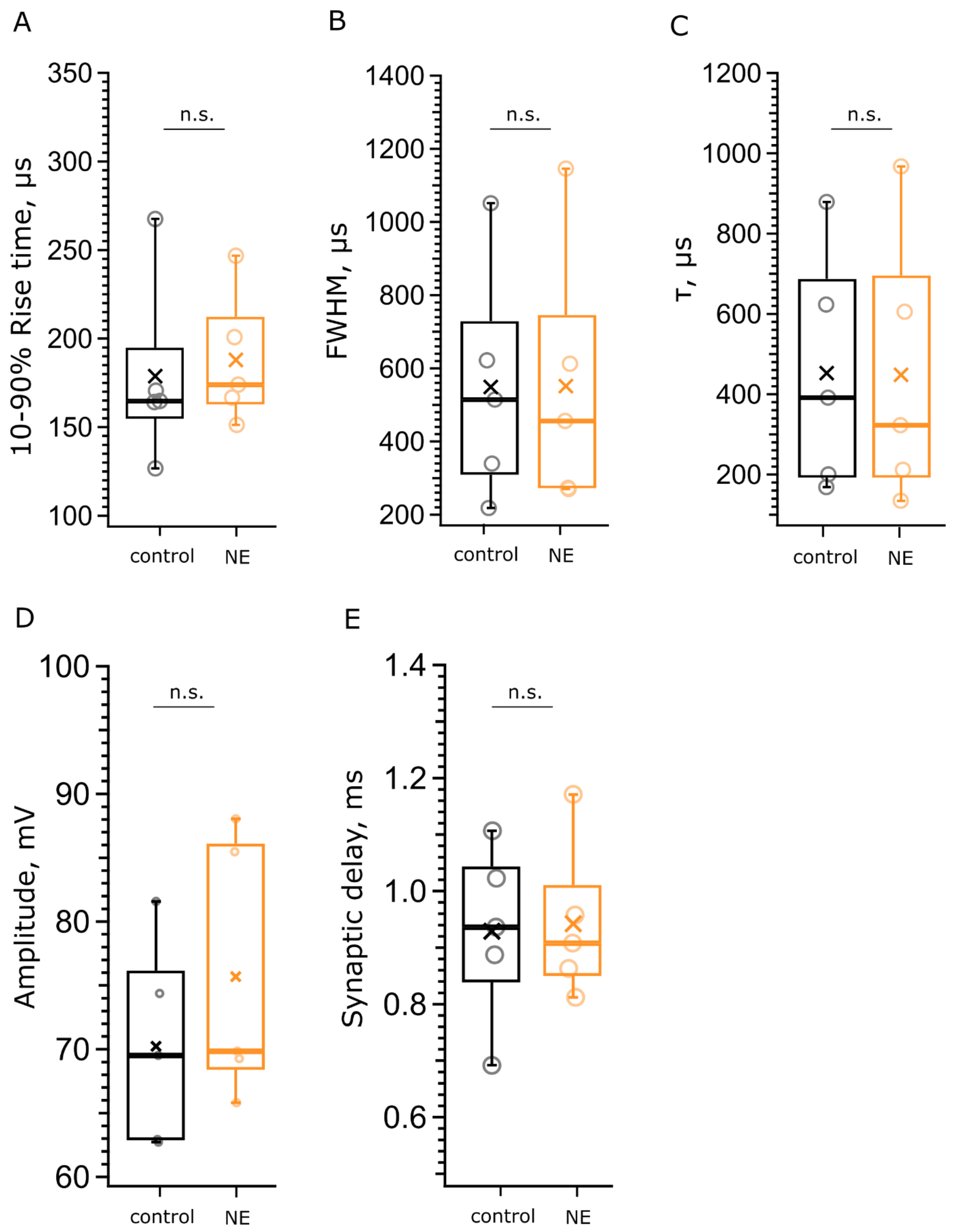


**Figure S4 Probing for effects of NE on the first AP at 100Hz stimulation frequency.** The acquisition protocol is described in Fig. S3. N= 5 cells from 4 mice. We first acquired AP trains in the control solution (black), followed by recording responses to the same sequence of stimuli, while bathing the calls with the 100 µM NE solution (orange). A, 10-90% rise time of the AP, *p=0.7590*, B – FWHM of the AP, p=0.9908, C, τ, *p=0.9849*, D, Synaptic delay, *p=0.8178*, E, Amplitude of the AP, *p=0.3705*. Each data point represents an average of the given parameter across the events in one mEPSC recording.

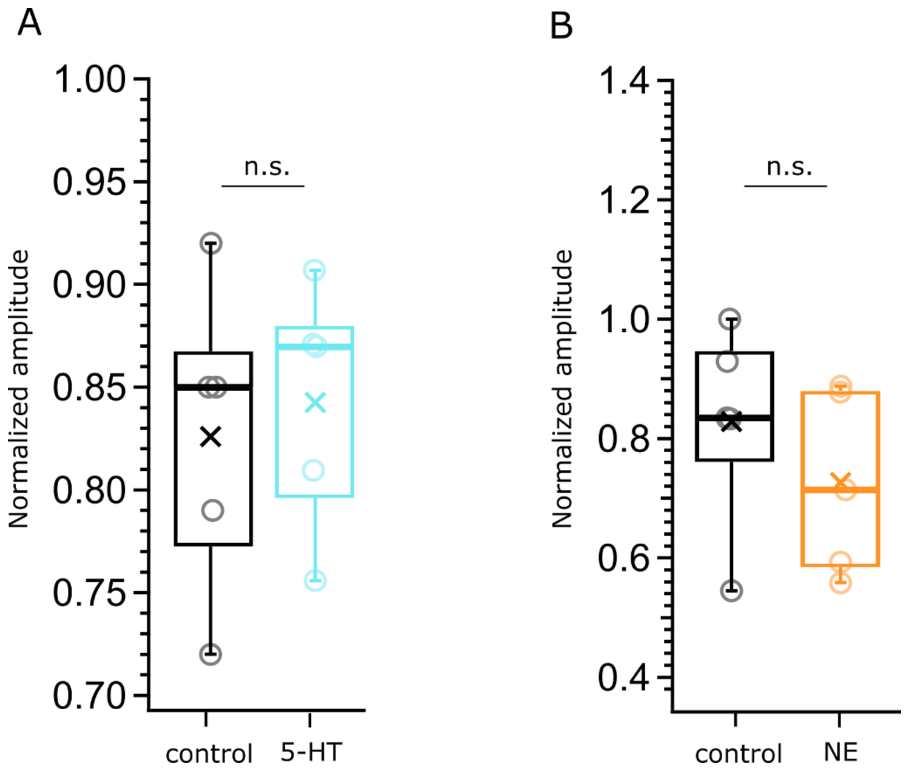


**Figure S5 Relative amplitude of the last AP of BC firing during 100 Hz train stimulation in control, 5-HT and NE conditions.** We normalized the amplitude of the 50th AP to that of the first AP of the train and compared the normalized amplitude between control – 5-HT (10 µM) and control – NE (100 µM). For both experiments we obtained consequential recordings from 5 cells each. A, Evaluating the effect of 5-HT on the amplitude of the last AP, control – black, 5-HT cyan, *p=0.7024*. B Evaluating the effect of NE on the amplitude of the last AP, control – black, NE – orange, *p=0.3532*

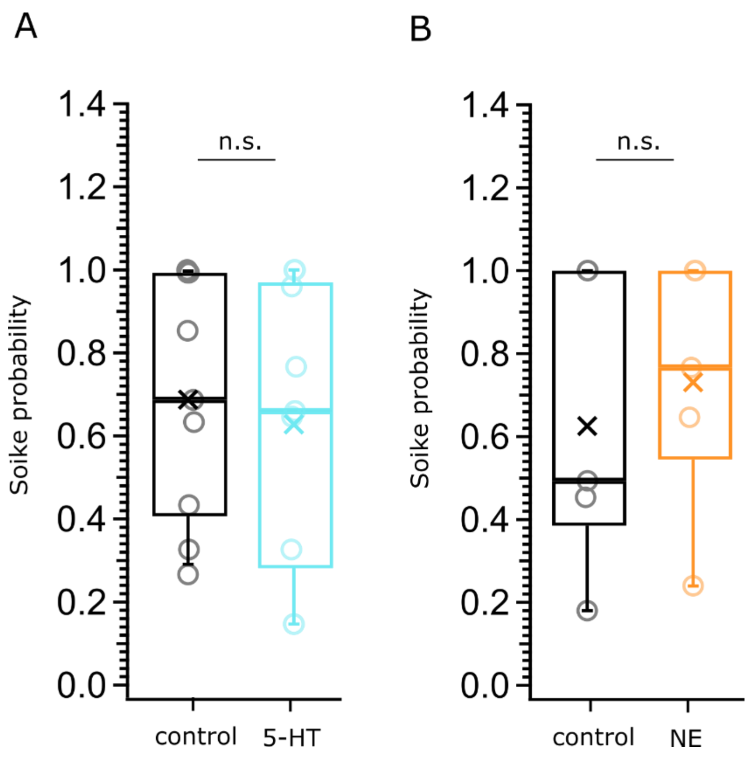


**Figure S6 BC spike probability at 200 Hz.** At lower frequencies the BCs that we studied did not display failures during the trains. APs were normalized by the 1 AP. The SP was calculated as the number of AP divided by the number of stimuli (50) that the afferent fibers were subjected to. A, Evaluating the effects of 5-HT on the BC SP, control in black, 5-HT in cyan, N=9 cells, *p=0.70*; B, Evaluating the effects of NE on the BC SP, control in black, NE in orange, N=5 cells, p=0.63. The data is represented as box plots with minimum, first quartile, median, third quartile, and maximum, the cross represents the mean. n.s. – non-significant
